# YY2/BUB3 axis-mediated SAC hyperactivity determines tumor cell fate through chromosomal instability

**DOI:** 10.1101/2023.10.07.561167

**Authors:** Rendy Hosea, Wei Duan, Ian Timothy Sembiring Meliala, Wenfang Li, Mankun Wei, Sharon Hillary, Hezhao Zhao, Makoto Miyagishi, Shourong Wu, Vivi Kasim

## Abstract

Spindle assembly checkpoint (SAC) is a crucial safeguard mechanism of mitosis fidelity, which is fundamental for equal division of duplicated chromosomes to the two progeny cells. Impaired SAC can lead to chromosomal instability (CIN), a well-recognized hallmark of cancer that facilitates tumor progression; paradoxically, high CIN levels are associated with better drug sensitivity and prognosis. However, the mechanism by which CIN determines tumor cell fates and drug sensitivity remain poorly understood. In this study, using a cross-omics approach, we identified YY2 as a mitotic regulator that peaks at M phase and promotes SAC activity by positively regulating the transcriptional activity of *budding uninhibited by benzimidazole 3* (*BUB3*), a component of SAC. While inducing CIN, YY2/SAC activity defect enhanced mitosis and tumor growth, whereas YY2/SAC hyperactivation, as a result of *YY2* overexpression, triggered mitotic delay and growth suppression. Furthermore, we revealed that excessive CIN, caused by either *YY2* overexpression or further inhibiting SAC activity in *YY2*-knocked out cells, leads to higher cell death rates. However, residual tumor cells that survived DNA damage-based therapy had moderate CIN and increased drug resistance; meanwhile *YY2* overexpression in these cells sensitizes them to DNA-damage agents. Hence, this study provides insights into the regulatory mechanism of SAC activity as well as the role of YY2/BUB3 axis, SAC activity, and CIN levels in determining tumor cell fate. Furthermore, this study also links up tumor cells drug resistance with moderate CIN, and suggest a novel anti-tumor therapeutic strategy that combines SAC activity modulators and DNA-damage agents.

**Significance:** This study identifies the novel role of YY2/BUB3 axis as a SAC modulator, as well as emphasizing the role of YY2-mediated SAC activity and CIN levels in determining tumor cell fates.

## Introduction

Chromosomal instability (CIN) is characterized by an increased rate of chromosomal structural and numerical abnormalities due to aberrant segregation. Chromosomal abnormalities cause significant fitness costs to normal cells, leading to various disorders, including metabolic alterations, proteotoxic stress, cell cycle arrest, and senescence (1,2). Furthermore, it is the most common cause of spontaneous abortion and severe developmental defects (3). Despite these deleterious effects, CIN is a well-recognized cancer hallmark, with approximately 90% of tumors displaying complex karyotypes, including structural and numerical CIN (4). Moreover, studies using mouse models support the notion that CIN significantly promotes tumor initiation (4). However, paradoxically, studies with human clinical tumor samples has revealed that high CIN levels are associated with improved prognosis and better responses to therapies, such as those using DNA-damage agents, thus demonstrating a tumor suppressive function (5-7).

Impaired activity of the spindle assembly checkpoint (SAC), one of the major cell cycle checkpoints, is a major cause of CIN (8). As a checkpoint at the metaphase–anaphase transition point, it ensures that sister chromatids are segregated equally into the two progeny cells and is thus the last checkpoint to guarantee mitotic fidelity through the passage of correct, intact genetic information to the progeny cells (9). SAC activity prevents premature cohesion cleavage, sister chromatid segregation, and mitotic exit by suppressing securin and cyclin B ubiquitination/proteasomal degradation before all sister chromatids are lined up at the equator and attached to spindle microtubules (9). Defect in SAC triggers premature sister chromatid segregation and mitotic exit, leading to increased CIN and faster cell proliferation, thereby serving as a driving force for tumorigenesis (10,11). Similarly, SAC hyperactivation also induces CIN; however, it leads to mitotic delay and tumor suppression (12,13). The role of SAC hyperactivation in tumorigenesis remains poorly understood. Furthermore, the reasons underlying the different outcomes related to SAC defects and hyperactivation remain to be explored.

Although mutations in SAC genes are rare, alterations in SAC gene expression are frequently found in tumor cells, suggesting a crucial role for transcriptional regulation in aberrant SAC activity in tumor cells (14,15). However, the mechanisms underlying SAC transcriptional regulation have not yet been fully elucidated. In this study, we utilized a cross-omics approach to search for potential transcriptional regulators of SAC genes and identified yin yang 2 (YY2) as a novel *budding uninhibited by benzimidazole 3* (*BUB3*) transcriptional regulator. Our results showed that YY2/BUB3 axis positively modulates SAC activity, prolongs mitotic time, and suppresses colorectal cancer (CRC) cells survival. However, despite of their opposite effects on SAC activity and CRC cells survival, both YY2/BUB3 deficiency and overexpression lead to increased CIN. We further revealed that different CIN degrees induced by YY2/BUB3 alterations is crucial for determining CRC cell fate, notably, excessive CIN level leads to cell death while moderate CIN is beneficial for CRC cells. Finally, our results showed a strong correlation between moderate CIN and drug resistance, such as that found in residual tumor cells resistant to DNA damage-inducing agents and/or SAC activation. In contrast, the induction of excessive CIN in those cells by *YY2* overexpression sensitizes them to DNA damage-inducing agents, suggesting a novel anti-tumor therapeutic approach by combining SAC activator and DNA damage-inducing agents.

## Materials and Methods

### Plasmids and constructs

shRNA expression vectors targeting two different sites of BUB3 or p31/comet were constructed using algorithm and method as described previously (16). Target sequences were predicted as follows: 5ʹ- GTC CAG AAG TGA ATG TGA T -3ʹ (shBUB31-1); 5ʹ- AGA AGA AGA AGT ATG CCT T -3ʹ (shBUB3-2); 5ʹ- GGC AGA CCT TCC ATC CGA A -3ʹ (shp31-1); 5ʹ- GGC AGA ACT GGA GAG TGT C -3ʹ (shp31-2).

*YY2* (NM_206923.4) overexpression vector (pcYY2), mutant *YY2* overexpression vectors, and lentivirus vector overexpressing *YY2* (pLenti-YY2) were constructed as described previously (17). For *BUB3* (NM_004725.4) overexpression vectors, human complementary DNA (cDNA) obtained by reverse-transcribing the total RNA extracted from HCT116 cells using the PrimeScript RT Reagent Kit with gDNA Eraser (Takara Bio, Dalian, China) was used as the template for amplifying the corresponding regions using the PrimeSTAR Max DNA Polymerase (Takara Bio). The amplicons were then cloned into the *Hind*III and *Eco*RI sites of pcEF9-Puro vector (18). For doxycycline-inducible *YY2* overexpression vector (pTRIPZ-YY2), *YY2* coding region was inserted into *Xho*I and *Mlu*I sites of tetracycline-inducible lentiviral pTRIPZ vector (GE Dharmacon technology, Lafayette, CO).

For reporter vectors bringing different regions of *BUB3* promoter (Refseq No. NC_000010.11; BUB3-Luc-1 with the −2,688 to +166 region; BUB3-Luc-2 with the −1,765 to +166 region; BUB3-Luc-3 with the −1,224 to +166 region; BUB3-Luc-4 with the −723 to +166 region; and BUB3-Luc-5 with the −658 to +166 region), the corresponding regions of the *BUB3* promoter were cloned into *Xho*I and *Hind*III sites of the pGL4.13 (Promega, Madison, WI). Human genome DNA extracted from HCT116 cells using the TIANamp Genomic DNA Kit (Tiangen Biotech, Beijing, China) was used as template for amplifying the promoter regions. BUB3-luciferase vector with mutated predicted YY2 binding site (BUB3-Luc^mut^) was constructed based on the site-specific mutagenesis method using a Site-directed Gene Mutagenesis Kit (Beyotime Biotechnology, Shanghai, China).

### Cell lines and cell culture

HCT116 (catalog number: TCHu 99) and HEK293T (catalog number: GNHu17) cells were purchased from the Cell Bank of Chinese Academy of Sciences (Shanghai, China) and cultured in Dulbecco’s modified Eagle’s medium (Gibco, Life Technologies, Grand Island, NY) with 10% FBS (Biological Industries, Beith Haemek, Israel) and 1% penicillin-streptomycin. Cell lines were verified using short-tandem repeat profiling method and were tested periodically for mycoplasma contamination using Mycoplasma Detection Kit-QuickTest (Biotool, Houston, TX). Transfection was performed using Lipofectamine 2000 (Invitrogen Life Technologies, Carlsbad, CA) according to the manufacturer’s instruction. HCT116^YY2KO^ cell with deletion of nucleotides located in +95 to +151 region (56 bp) of *YY2* coding sequence was established using CRISPR/Cas9 method as described previously.(17) For transfection, cells were seeded in 6-well plates and transfected with 2 µg of indicated shRNA expression vectors and/or overexpression vectors. Twenty-four hours after transfection, puromycin selection (final concentration: 1 μg/ml) was performed for 36 h to eliminate untransfected cells.

Lentiviruses for establishing *YY2*-inducible cell lines and stable cell lines for xenograft experiments were generated by co-transfecting HEK293T cells with 8 μg pLenti-YY2 or pTRIPZ-YY2 vectors, 6 μg pCMVΔR, and 2 μg pCMV-VSVG in a 10 cm dish. Growth media was changed the following day and lentivirus-containing supernatant was harvested and filtered with a 0.45-μm filter after 48 h. For infection, HCT116 cells were cultured in 6-well plates. Twenty-four hours later, the medium was changed to 1 ml fresh culture medium and 1 ml corresponding lentivirus supernatant. Infected cells were then selected using 1 μg/ml puromycin for 7 days.

For oxaliplatin treatment, cells were seeded in 6-well plates and cultured with medium containing indicated concentration of oxaliplatin (MedChemExpress, Monmouth Junction, NJ).

### Clinical human colon carcinoma specimen

Human colon carcinoma specimens were obtained from colon carcinoma patients undergoing surgery at Chongqing University Cancer Hospital (Chongqing, China), and stored in Biological Specimen Bank of Chongqing University Cancer Hospital. Patients did not receive chemotherapy, radiotherapy, or other adjuvant therapies prior to the surgery. The specimens were snap-frozen in liquid nitrogen. Prior patient’s written informed consents were obtained. The experiments were approved by the Institutional Research Ethics Committee of Chongqing University Cancer Hospital, and conducted in accordance with Declaration of Helsinki.

### Animal experiments

BALB/*c-nu*/*nu* mice (Strain number D000521, male, body weight: 18–22 g, 6 weeks old) were purchased from the Chongqing Medical University (Chongqing, China), randomly divided into 4 groups (n = 7), and injected subcutaneously with 3 × 10^6^ indicated stable cell lines. Oxaliplatin (MedChemExpress) was administered intraperitoneally at a dose of 5 mg/kg twice a week for 3 weeks. Treatment began on day 10, when the tumor volume reached 100 mm^3^. Tumor size (V) was evaluated by a caliper every 2 days using the following equation: V = a × b^2^/2, where a and b are the major and minor axes of the tumor, respectively. The investigator was blinded to the group allocation and during the assessment. Animal studies were approved by the Institutional Ethics Committee of Chongqing Medical University, and carried out in the Chongqing Medical University. All animal experiments conformed to the approved Guidelines for the Care and Use of Laboratory Animals of Chongqing Medical University. All efforts were made to minimize suffering.

### Metaphase spread

Cells were prepared as described above and arrested at metaphase by 2 μg/ml colchicine (Aladdin, Shanghai, China) treatment for 3 h before being harvested, re-suspended in hypotonic solution (0.075 M KCl) and incubated for 15 min at 37 °C. After cell swelling, cells were fixed using 2 ml of freshly prepared methanol-acetic acid fixative (3:1) and collected by centrifugation. The pellet was re-suspended in 5 mL of 3:1 methanol-acetic acid for 30 min and dropped onto pre-cooled slides. Images of mitotic chromosomes were acquired with fluorescence microscope (Olympus BX53). Chromosome number per cell was quantified.

### Karyotype and G-band analysis

Cells were treated with 2 μg/ml colchicine for 1 h and collected processed as described above. After dropped onto pre-cooled slides, the slides were allowed to dry overnight at room temperature, banded with trypsin and stained with 10% Giemsa stain (Biosharp, Shanghai, China). Banded metaphases were photographed using light microscopy with Olympus BX53 microscope and analyzed using SmartType software (Digital Scientific, Cambridge, UK). 10 metaphase spread images were analyzed for each sample.

### Cell cycle enrichment and DNA content analysis

For cell cycle analysis, cells were prepared as described above and synchronized in G_0_ phase by FBS starvation for 24 h. Cells were washed two times in phosphate-buffered saline (PBS) and either collected immediately (0 h) or further cultured in medium with 10% FBS for indicated time points. Percentages of each cell cycle phases were estimated using ModFit LT (Verity Software House, Topsham, ME).

For DNA content analysis, cells were prepared as described above and subjected to flow cytometry after staining the DNA with propidium iodide (PI) (NeoBiosciences). Percentage of cells with each DNA content (2N, 3N, 4N, 5-8N) were estimated using CytExpert software (Beckman Coulter). For DNA content analysis using sorted cells, cells were transfected or infected with indicated vectors and sorted for living and dying cells as described above before being stained with PI.

### Chromosome bridge and micronucleus staining

Cells were prepared as described above and seeded in 3.5-cm confocal dishes (3 × 10^4^ cells per dish), incubated overnight, and synchronized at prometaphase by nocodazole treatment (final concentration: 100 ng/ml; Beyotime Biotechnology) for 6 h. Cells were then fixed with 4% paraformaldehyde for 30 min at room temperature and permeabilized with PBS containing 0.1% Triton X-100 for 5 min. Nuclei were stained with DAPI (Beyotime Biotechnology) for 15 min. For chromosome bridge observation, images were taken using laser scanning confocal microscopy (Leica Microsystems TCS SP5, Heidelberg, Germany), and quantification was performed by LAS X software (Leica Microsystems). For micronuclei observation, images were taken using fluorescence microscope (Olympus IX71). Micronucleus rate was defined as the ratio of total micronuclei to total cell number.

For chromosome bridge observation in xenografted tumor lesions, fresh xenografted tumor lesions were fixed using 4 % paraformaldehyde for overnight, embedded in paraffin, and sectioned at 4 μm thickness using a cryostat. After being dewaxed using xylene and rehydrated, sections were stained with 0.5% hematoxylin-eosin (Sangon Bio, Shanghai, China). After samples were dehydrated and mounted in coverslip, images were taken using light microscope (Olympus BX53).

For micronucleus rate assay using sorted cells, cells were prepared and sorted for living and dying cells as described above. Cells were then fixed on slides before being fixed, permeabilized, and stained as described above. Quantification of nucleus size was performed using ImageJ.

### Live-cell imaging

For fluorescence imaging of mitotic division, cells were prepared as described above prior to seeding in 3.5-cm glass bottom dish (Wuxi NEST Biotechnology, Wuxi, China; 1 × 10^5^ cells/dish) and grown overnight. Nuclei was stained using Hoechst 33342 (Solarbio, Beijing, China) 30 min prior to imaging. The culture was maintained at 37 °C under 5% CO_2_ in a stage-top incubator (Tokai Hit, Shizuoka, Japan). Images were acquired every 5 min for 6 h with a 20-ms exposure time using inverted fluorescence microscope Olympus IX83 (Olympus, Tokyo, Japan). Images were processed using FLUOVIEW (v.2.3). Mitotic time was quantified as the time from nuclear envelope breakdown (NEBD) until the onset of anaphase. To study cumulative mitotic exit, nocodazole (final concentration: 100 ng/ml; Beyotime Biotechnology) was added, and the cells were imaged every 10 min for 10 h using Olympus IX83 microscope. The number of cells that exited mitosis was quantified and analyzed over time using FLUOVIEW software.

### RNA extraction, quantitative real-time PCR (qRT-PCR), and western blotting

Detailed methods for performing western blotting and quantitative PCR analysis are described in the Supplementary Materials and Methods. The sequences of the primers and antibodies used are shown in Supplementary Tables S1 and S2, respectively. Images of uncropped blots are shown in Supplementary Fig. S12A to S12K.

### Statistical analysis

All values of the experimental results were presented as mean ± SD (n = 3; unless further indicated). Quantification data were analyzed by one-way ANOVA conducted using SPSS Statistics (v. 20.0.) A value of *P* < 0.05 was considered statistically significant. For each experiment the number of technical replicates, and the number of independent experiments in each group is reported in Figure legends.

## Results

### Identification of transcriptional regulators modulating SAC

To identify a novel modulator of the SAC and subsequently tumor cell CIN, we screened genes involved in both “mitotic sister chromatid segregation” and “mitotic spindle assembly checkpoint signaling” from the Gene Ontology (GO) database (QuickGO, http://www.ebi.ac.uk/QuickGO; GO numbers: 0000070 and 0007094; respectively) and obtained 195 genes. After excluding non-human-origin and hypothetical genes, we further screened genes that have been reviewed and annotated using the manual-curation process in UniProtKB and referenced based on a PubMed Unique Identifier (PMID). The 20 genes identified in this screening were subjected to transcription factor (TF) enrichment analysis using enrichment analysis version 3 (ChEA3, https://maayanlab.cloud/chea3/), a web server application developed to conduct TF enrichment analysis utilizing gene set libraries from published omics assay data extracted from multiple sources (19) (Fig. 1A). Among the top 17 TFs obtained, eight TFs were known SAC modulators, thus confirming the validity of this screening method (20-23). whereas nine TFs were potential novel SAC modulators (Fig. 1B).

**Figure 1.**
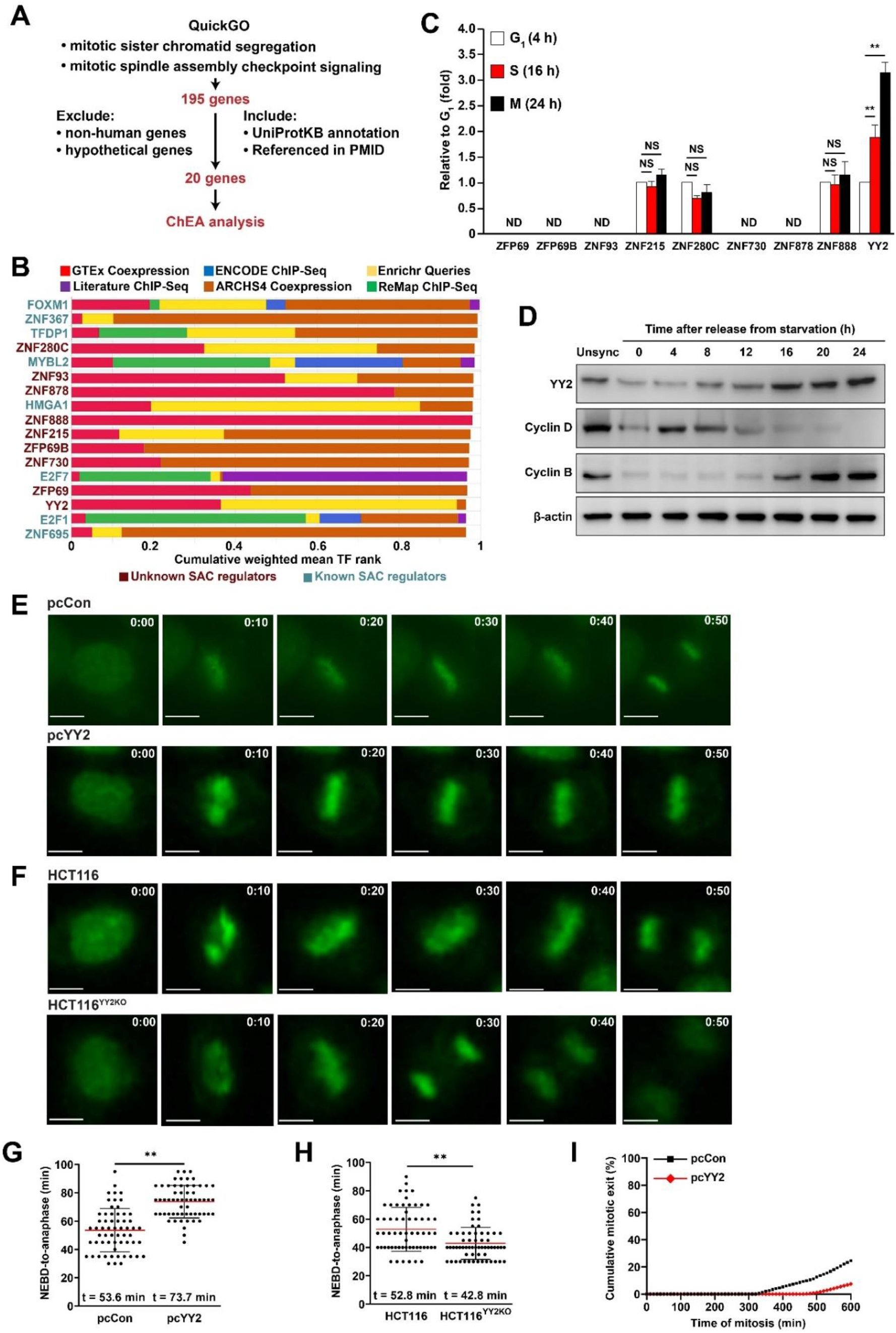
Identification of novel transcriptional regulators modulating SAC. **A,** Schematic diagram of screening strategy for potential novel transcriptional regulators modulating SAC. **B,** Top 17 potential TFs modulating SAC obtained from ChEA analysis. **C,** mRNA expression levels of potential novel SAC modulators at each cycle phase, as examined using qRT-PCR. **D,** YY2, cyclin D, and cyclin B protein expression levels in HCT116 cells at indicated time-points after serum starvation release, as determined by western blotting. **E–F,** Mitotic time of *YY2*-overexpressed HCT116 cells (**E**) and HCT116^YY2KO^ cells (**F**), as determined using time-lapse microscopy. Representative images (scale bars: 20 μm) are shown. **G–H,** Scatter plots showing the time-length from nuclear envelope breakdown (NEBD) to anaphase of *YY2*-overexpressed HCT116 cells (**G**) and HCT116^YY2KO^ cells (**H**) (n = 60, pooled from three independent experiments). **I,** Time-lapse analysis of duration of mitosis in *YY2*-overexpressed HCT116 cells arrested with nocodazole (final concentration: 0.2 μM; n = 150, pooled from three independent experiments). Cells transfected with pcCon, wild-type HCT116, or unsynchronized cells were used as controls. β-actin was used for qRT-PCR normalization and as western blotting loading control. Quantification data are shown as mean ± SD. All data were obtained from three independent experiments. *P* values were calculated by one-way analysis of variance (ANOVA). pcCon: pcEF9-Puro; Unsync: unsynchronized; \*\**P* < 0.01; NS: not significant; ND: not detected.

TFs that regulate the cell cycle usually exhibit oscillating levels during cell cycle progression (24). Thus, we first determined the timing of each cell cycle phase by synchronizing HCT116 CRC cells in the G_0_/G_1_ phase using serum starvation (Supplementary Fig. S1A). The percentage of cells in the G_0_/G_1_ phase decreased from 78.12% immediately after serum starvation release to 14.11% 16 h later, whereas cells in the S phase increased from 10.20% to a peak at 76.89% after 16 h, and cells in the G_2_/M phase reached 55.69% at 24 h after serum starvation release. Cell cycle-dependent expression analysis of the nine novel SAC modulators revealed that significant expression of ZFP69, ZFP69B, ZNF93, ZNF730, and ZNF878 could not be detected in any cell cycle state and that of ZNF215, ZNF280C, and ZNF888 did not show significant differences during cell cycle progression. Meanwhile, YY2 mRNA expression clearly increased during cell cycle progression and peaked during the M phase (Fig. 1C). A more detailed time-course investigation showed that YY2 mRNA expression peaked at 20 h (Supplementary Fig. S1B), whereas its protein level started to increase significantly at 16 h and peaked at 24 h after serum starvation release (Fig. 1D). Meanwhile, the levels of G_1_ and G_2_/M phase cyclins, specifically cyclins D and B, peaked at 4 h and 24 h after serum starvation release, respectively, further confirming the relationship between YY2 expression and the M phase.

SAC delays mitotic exit by stabilizing securin and suppressing APC/C activity. To further explore the role of YY2 in regulating SAC activity and subsequently mitotic exit, we altered *YY2* expression in HCT116 cells, a near-diploid human CRC cell line with a mitotic time similar to that of normal cells and considered CIN-negative cells (Supplementary Fig. S1C– S1D) (25). As shown by time-lapse microscopic images, *YY2* overexpression delayed the mitosis, whereas *YY2* knockout significantly accelerated it (Fig. 1E–1F, Supplementary Videos S1–S4). Measurement of mitotic time revealed that *YY2* overexpression prolonged the mitotic time from 53.6 min to 73.7 min, while *YY2* knockout significantly shortened it from 52.8 min to 42.8 min (Fig. 1G–1H). Next, we modulated *YY2* expression and synchronized cells in the M phase using nocodazole. *YY2* overexpression significantly slowed down the degradation of securin and cyclin B, an event that marks mitotic exit (Supplementary Fig. S2A–S2B), whereas knocking out *YY2* accelerated their degradation (Supplementary Fig. S2C–S2D). Moreover, to validate *YY2* overexpression impact on SAC activity, we investigated the ability of *YY2*-overexpressed cells treated with nocodazole, a metaphase-arresting SAC activator, to sustain an extended mitotic arrest (26). *YY2* overexpression led to stronger SAC activity, as demonstrated by fewer cells that exit mitosis under constant SAC activation; thereby solidifying the role of YY2 in promoting SAC activity (Fig. 1I). These results revealed that YY2, of which the expression oscillates during the cell cycle and reaches its peak in the M phase, is a novel SAC modulator that controls mitotic progression.

### *YY2-*mediated SAC regulation is crucial for its tumor suppressive effect

YY2 is a zinc-finger protein and has been reported to have a tumor suppressive function (17,27-29). A comparative analysis using clinical CRC tissues and corresponding normal adjacent tissues revealed a significant decrease in YY2 mRNA and protein levels in CRC tissues (Supplementary Fig. S3A–S3B). Furthermore, *YY2* overexpression significantly suppressed the viability of HCT116 cells (Supplementary Fig. S3C). As a decrease in cell viability might be due to a decrease in proliferation, increase in cell death, or both, we next examined the effect of YY2 on HCT116 cell proliferation and cell death. *YY2* overexpression robustly suppressed CRC cell proliferation, as indicated by the decrease in EdU-positive cells and colony-formation potential (Supplementary Fig. S3D–S3E), while increasing the cell death rate (Fig. 2A), suggesting that YY2 may exert its tumor suppressive function most plausibly by decreasing cell proliferation and promoting cell death. In contrast, the viability, proliferation, and colony-formation potential of HCT116^YY2KO^ cells were significantly higher than those of wild-type cells (Supplementary Fig. S3F–S3H). Interestingly, *YY2* knockout did not significantly affect the cell death rate (Fig. 2B), most plausibly because of the overall low cell death rate.

**Figure 2.**
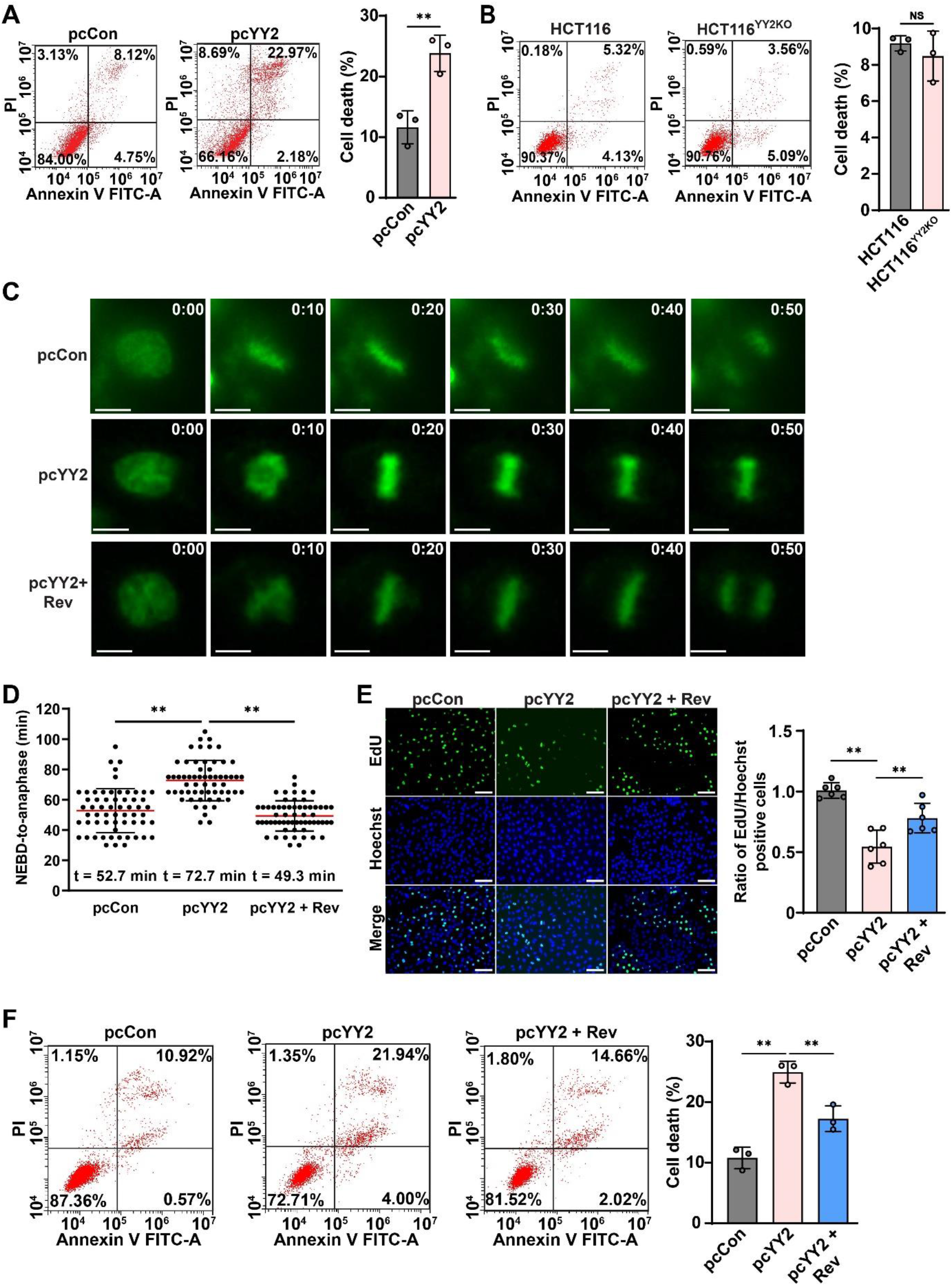
YY2 suppresses tumorigenesis by activating SAC activity. **A–B,** Cell death rate of *YY2*-overexpressed HCT116 (A) and HCT116^YY2KO^ cells (B), as examined using Annexin V/PI staining and flow cytometry. **C–D,** Mitotic time of *YY2*-overexpressed, reversine-treated HCT116 cells, as determined using time-lapse microscopy. Representative images (**C**; scale bars: 20 μm) and scatter plot showing the time-length from NEBD to anaphase (**D**; n = 60, pooled from three independent experiments) are shown. **E,** Proliferation potential of *YY2*-overexpressed, reversine-treated HCT116 cells, as determined using EdU-incorporation assay. Representative images (left; scale bars: 200 μm) and ratio of proliferative cells (right; each dot represents the mean value of three technical replicates) are shown. **F,** Cell death rate of *YY2*-overexpressed, reversine-treated HCT116 cells, as examined using Annexin V/PI staining. Cells transfected with pcCon or wild-type HCT116 cells were used as controls. Quantification data are shown as mean ± SD. All data were obtained from three independent experiments. *P* values were calculated by one-way ANOVA. pcCon: pcEF9-Puro; Rev: reversine (final concentration: 0.2 μM; \*\**P* < 0.01; NS: not significant.

To examine whether YY2 exerts its tumor suppressive effect by regulating SAC activity, we treated *YY2*-overexpressed HCT116 cells with reversine, a SAC activity inhibitor. Reversine treatment reduced cyclin B and securin accumulation induced by *YY2* overexpression (Supplementary Fig. S3I) and re-suppressed mitotic time prolonged by *YY2* overexpression from 72.7 min to 49.3 min (Fig. 2C and 2D, Supplementary Video S5). Accordingly, reversine treatment restored the proliferation, viability, and colony-formation potential of *YY2*-overexpressed HCT116 cells (Fig. 2E; Supplementary Fig. S3J–S3K) while suppressing *YY2* overexpression-induced cell death (Fig. 2F). These results reveal that YY2-mediated SAC activation and mitotic regulation are crucial for exerting its tumor suppressive effect.

### YY2 regulates SAC by directly activating *BUB3* transcription

To analyze the molecular mechanism underlying YY2-mediated regulation of SAC activity, we performed a cross-omics analysis to identify its potential transcriptional target. To this end, RNA-sequencing data obtained in our previous study (https://www.ncbi.nlm.nih.gov/geo/query/acc.cgi?acc=GSE184138) (17) using *YY2*-overexpressed HCT116 cells were analyzed for differentially expressed genes (RNA-seq; Fig 3A). The differentially expressed genes (fold-change > 1.01; *P* < 0.05) were then analyzed for enriched GO terms and Reactome pathways using the Database for Annotation, Visualization, and Integrated Discovery (DAVID, https://david.ncifcrf.gov/). GO analysis indicated that *YY2* overexpression enriched genes involved in cell division, mitotic nuclear division, sister chromatid cohesion, and negative regulation of ubiquitin-protein ligase activity in mitosis (Fig. 3B); meanwhile, Reactome analysis revealed the enrichment of APC/C-Cdc20-mediated degradation of securin, separation of sister chromatids, mitotic prometaphase, and resolution of sister chromatid cohesion (Fig. 3C), indicating a possible prominent role of YY2 in regulating sister chromatid segregation.

**Figure 3.**
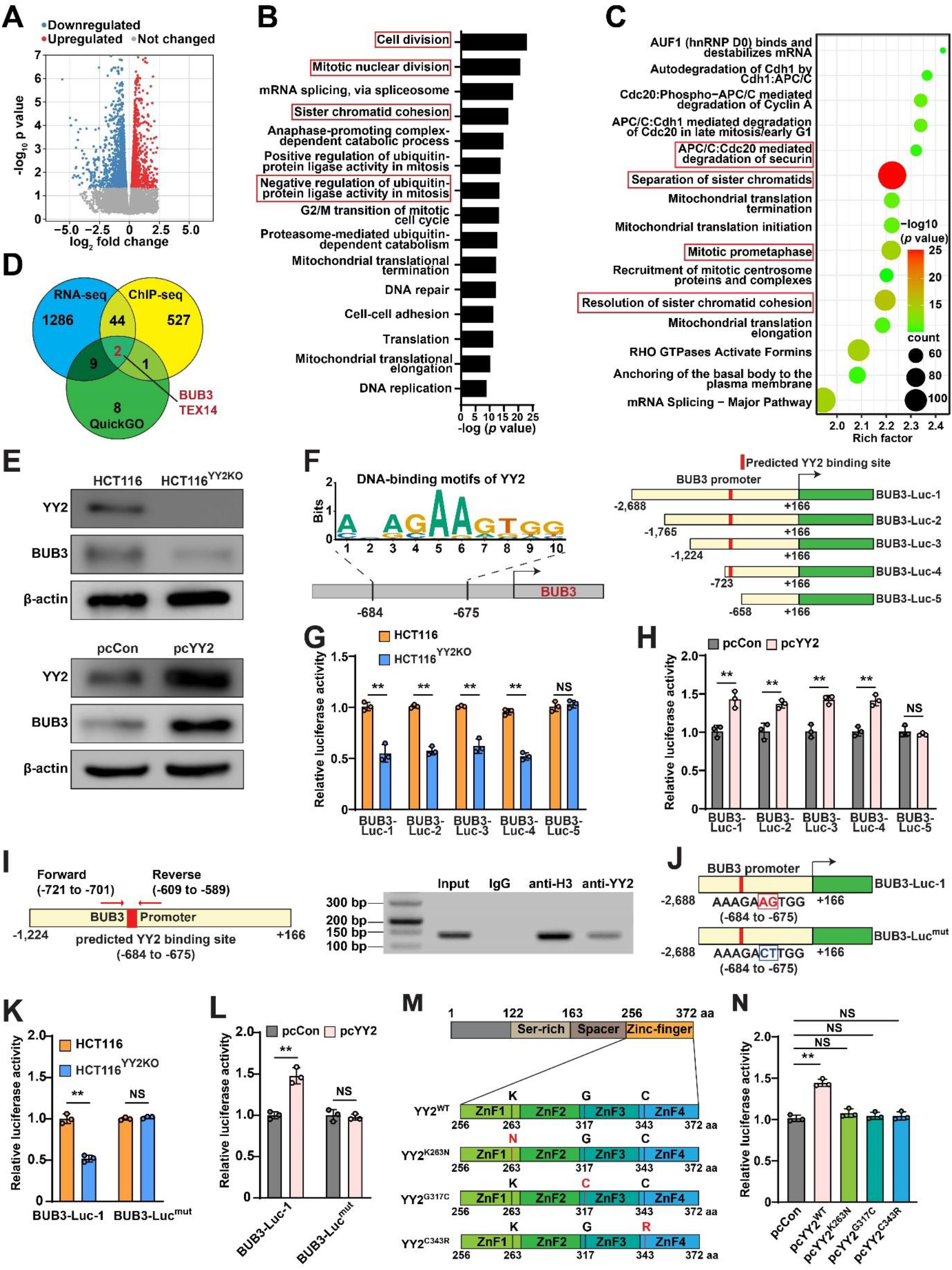
YY2 directly regulates *BUB3* transcription. **A,** Volcano plot of log2 fold-change versus adjusted *P* value for gene expression changes (fold-change > 1.01-fold; *P* value < 0.05) in *YY2*-overexpressed HCT116 cells, as analyzed by RNA-seq. **B–C,** GO (**B**) and Reactome (**C**) enrichments of differentially expressed genes in *YY2*-overxpressed HCT116 cells. **D,** Overlapping genes upregulated by YY2 based on RNA-seq, predicted YY2 target genes identified by ChIP-seq, and QuickGO screening. **E,** BUB3 protein level in *YY2*-overexpressed HCT116 and HCT116^YY2KO^ cells, as determined using western blotting. **F–H,** Schematic diagram (**F**) as well as relative luciferase activities of *BUB3* reporter vectors in HCT116^YY2KO^ cells (**G**) and *YY2*-overexpressed HCT116 cells (**H**). **I,** Binding capacity of YY2 to the predicted region in the *BUB3* promoter, as determined using ChIP assay. Location of the primer pair used for PCR are shown. **J–L,** Schematic diagram (**J**) as well as relative luciferase activities of BUB3-Luc^mut^ in HCT116^YY2KO^ cells (**K**) and *YY2*-overexpressed HCT116 cells (**L**). **M–N,** Schematic diagram of *YY2* mutants overexpressing vectors (**M**) and relative luciferase activity of BUB3-Luc-1 in HCT116 cells overexpressing indicated *YY2* mutants (**N**). Cells transfected with pcCon or wild-type HCT116 cells were used as controls. β-actin was used as western blotting loading control. Quantification data are shown as mean ± SD. All data were obtained from three independent experiments. *P* values were calculated by one-way ANOVA. pcCon: pcEF9-Puro; \*\**P* < 0.01; NS: not significant.

To identify potential YY2 transcriptional targets that regulate SAC activity, we overlapped differentially expressed genes identified by RNA-seq with those identified by chromatin immunoprecipitation (ChIP)-sequencing using an anti-YY2 antibody reported previously (27), as well as genes obtained from QuickGO screening (Fig. 1A). Two genes, *BUB3* and *TEX14*, were identified as potential YY2 direct transcriptional targets (Fig. 3D). BUB3 is a component of the SAC complex that regulates sister chromatid segregation and mitotic progression (9); meanwhile, TEX14 is required for the formation of intercellular bridges during meiosis (30). In accordance with previous studies (30,31), analysis of the TEX14 expression profiles using GTEx v8 expression data across 29 tissues revealed its testis specificity (Supplementary Fig. S4A). Furthermore, absolute qRT-PCR results showed that the copy number of *TEX14* was very low in HCT116 cells (Supplementary Fig. S4B). Meanwhile, BUB3 mRNA increased nearly two-fold upon *YY2* overexpression (Supplementary Fig. S4C), whereas its protein level was positively correlated with YY2 in *YY2*-knock-down and *YY2*-overexpressed HCT116 cells (Fig. 3E). Moreover, to avoid the effect of YY2 oscillation during cell cycle, we arrested the HCT116 cells at G_0_/G_1_ phase using serum starvation and investigated the effect of altering YY2 expression on BUB3. The results consistently demonstrated that *YY2* alteration positively correlates with BUB3 expression level, further confirming the direct regulation of YY2 on BUB3 expression (Supplementary Fig. S4D−S4E).

Next, we analyzed the *BUB3* promoter using JASPAR (http://jaspar.genereg.net/) (32), and identified a potential YY2-binding site at −684 to −675 of its promoter region (Fig. 3F). To confirm the role of this binding site in YY2-mediated regulation of *BUB3* transcriptional activity, we constructed reporter vectors comprising the −2,688 to +166 region (BUB3-Luc-1), −1,765 to +166 region (BUB3-Luc-2), −1,224 to +166 region (BUB3-Luc-3), −723 to +166 region (BUB3-Luc-4), and −658 to +166 region (BUB3-Luc-5) of the *BUB3* promoter (Fig. 3F). *YY2* knockout robustly suppressed the transcriptional activities of BUB3-Luc-1, BUB3-Luc-2, BUB3-Luc-3, and BUB3-Luc4, which contained the predicted YY2-binding site, but not that of BUB3-Luc-5, which lacked the predicted binding site (Fig. 3G). Concomitantly, *YY2* overexpression promoted the activities of BUB3-Luc-1, BUB3-Luc-2, BUB3-Luc-3, and BUB3-Luc4, but not that of BUB3-Luc-5 (Fig. 3H). These results suggest that the −723 to −659 region of the *BUB3* promoter is crucial for YY2-mediated regulation of its transcriptional activity. Furthermore, the result of a ChIP assay using an anti-YY2 antibody revealed that YY2 could bind to the −721 to −589 region of the *BUB3* promoter (Fig. 3I). Subsequently, mutations at the predicted YY2-binding site on the *BUB3* promoter (AAAGA*AG*TGG to AAAGA*CT*TGG; BUB3-Luc^mut^, Fig. 3J) abolished the YY2-mediated regulation of its transcriptional activity (Fig. 3K–3L). To further confirm YY2 transcriptional regulator activity on *BUB3*, we used mutant *YY2* overexpression vectors with a zinc finger mutation constructed previously based on cancer genomics data set from the cBioportal database (Fig. 3M) (17,33). Our results showed that these mutant YY2 failed to regulate *BUB3* promoter transcriptional activity (Fig. 3N). Together, these results showed that YY2 is a transcriptional regulator of *BUB3* that binds to the −684 to −675 site of its promoter region.

To elucidate the role of BUB3 in YY2-induced SAC activity and prolonged mitosis, we first analyzed the effect of *BUB3* overexpression on mitotic time. Similar to that of *YY2* overexpression, *BUB3* overexpression in HCT116 cells prolonged the mitotic time from 51.3 min to 80.5 min (Supplementary Fig. S4F–S4H and Supplementary Video S6) and slowed the degradation rates of cyclin B and securin proteins (Supplementary Fig. S4I–S4K). *BUB3* overexpression restored the length of HCT116 cells’ mitotic time, which was shortened to 43.3 min by *YY2* knockout, to a level similar to that observed in the control (50 min and 52.9 min, respectively; Fig S5A–S5C and Supplementary Video S7). Next, we constructed two shRNA expression vectors targeting different sites of BUB3 and selected shBUB3-2 (refers as shBUB3 hereafter), which had a higher suppressive effect, for further experiments (Supplementary Fig. S5D–S5E). Similar to the trend observed after knocking out *YY2*, knocking down *BUB3* shortened the mitotic time of HCT116 cells from 52.7 min to 40.5 min and resuppressed the mitotic time, which was prolonged by *YY2* overexpression, from 71.4 min to 46.8 min (Supplementary Fig. S5F–S5H and Supplementary Video S8–S9). Accordingly, *BUB3* knockdown restored HCT116 cell viability and cell death, which was suppressed by *YY2* overexpression (Supplementary Fig. S5I–S5J). Moreover, *BUB3* overexpression canceled the increase of CRC cell viability in *YY2* knockout HCT116 cells (Supplementary Fig. S5K). These results confirmed that YY2 regulates SAC and subsequently, mitotic progression and tumorigenic potential by enhancing *BUB3* expression.

### Both *YY2* knockout and overexpression induce CIN

Defects in SAC activity disrupt the mechanism that guarantees the proper arrangement of chromosomes at the equator and attachment to the mitotic spindle, making them prone to missegregation, and therefore, is a well-known causal factor of CIN induction. Thus, we examined the effect of *YY2* silencing on CIN indicators. We observed significant increases in the frequency of cells with DNA content > 4N (Fig. 4A) as well as in nuclear size (Fig. 4B) in HCT116^YY2KO^ cells, indicating the increase of the frequency of polyploid cells. Metaphase spread analysis results showed a significant increase in the karyotypic heterogeneity of HCT116^YY2KO^ cells, as nearly 80% of HCT116 cells had 45–47 chromosomes and fewer than 4% had more than 47 chromosomes, whereas only approximately 50% of HCT116^YY2KO^ cells had 45–47 chromosomes and the percentage of cells with more than 47 chromosomes increased to nearly 35% (Fig. 4C). Single karyotype analysis further confirmed the increase of cells with trisomy as well as karyotypic heterogeneity in HCT116^YY2KO^ (Fig. 4D).

**Figure 4.**
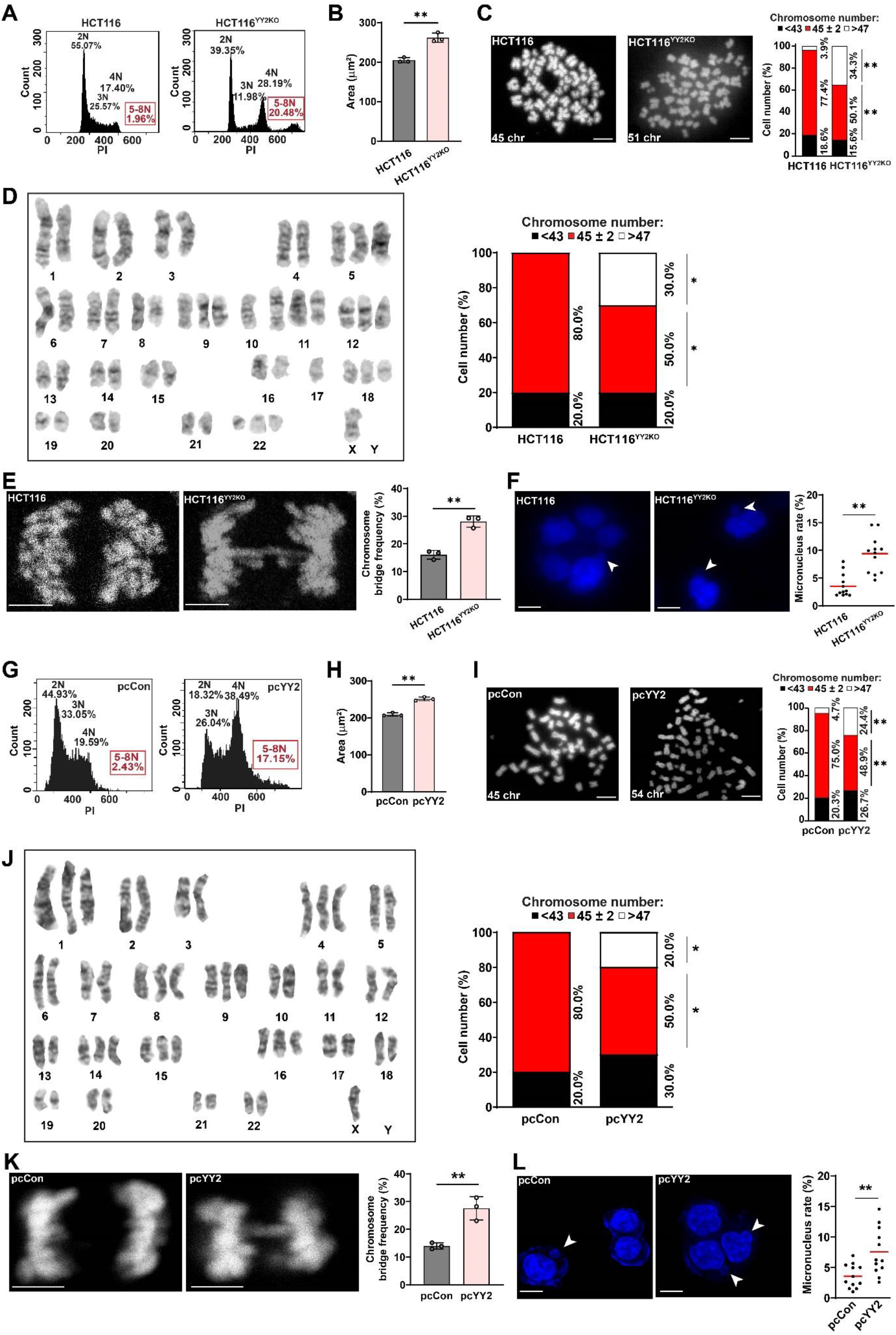
Both *YY2* knockout and overexpression induces CIN. **A,** DNA content in HCT116^YY2KO^ cells, as analyzed using PI staining and flow cytometry. **B,** Average nuclei size of DAPI-stained HCT116^YY2KO^ cells (each dot represents average nuclei size from one independent experiment, with total 100 cells/group). **C,** Chromosome number per cell in HCT116^YY2KO^ cells, as analyzed using metaphase spread. Representative images (scale bars: 10 μm) and percentage of cells with indicated chromosome number (total cells counted: 50 cells/group, pooled from three independent experiments) are shown. **D,** Single karyotype analysis of HCT116^YY2KO^ cells. Representative images and percentage of cells with indicated chromosome number (total cells counted: 10 cells/group, pooled from three independent experiments) are shown. **E,** Chromosome bridge frequency in HCT116^YY2KO^ cells. Representative images (scale bars: 5 μm) and quantification results (each dot represents chromosome bridge frequency from one independent experiment, with total 100 mitotic-cells/group) are shown. **F,** Micronucleus rate in HCT116^YY2KO^ cells. Representative images of micronuclei (indicated by arrowheads; scale bars: 20 μm) and micronucleus rate (ratio of micronuclei number to total cell number; each dot represents micronucleus rate/slide with > 100 cells/slides; four technical replicates from three independent experiments) are shown. **G– L,** DNA content analysis (**G**), average nuclei size (**H**), chromosome number per cell (**I**), single karyotype analysis (**J**), chromosome bridge frequency (**K**), and micronucleus rate (**L**) in *YY2*-overexpressed HCT116 cells. Cells transfected with pcCon or wild-type HCT116 cells were used as controls. Quantification data are shown as mean ± SD. All data were obtained from three independent experiments. *P* values were calculated by one-way ANOVA. pcCon: pcEF9-Puro; \**P* < 0.05; \*\**P* < 0.01.

Failed sister chromatid segregation leads to the formation of partly unsegregated sister chromatids or chromosome bridges, of which DNA fragments are subsequently wrapped by a nuclear membrane-like structure during cytokinesis to form a micronucleus, a small, nucleus-like structure in the cytoplasm. *YY2* knockout significantly increased the chromosome bridge frequency (Fig. 4E) and concomitantly increased the micronucleus rate (Fig. 4F), which could lead to the formation of aneuploid cell. Meanwhile, *BUB3* overexpression could partially abrogate the frequencies of polyploid cells and chromosome bridges, as well as the micronucleus rate in HCT116^YY2KO^ cells (Supplementary Fig. S6A–S6C). Together, these results suggest that defects in the YY2/BUB3 pathway could contribute to CIN induction in tumor cells, most plausibly due to impaired SAC activity.

Surprisingly, *YY2* overexpression, which induced SAC hyperactivation and mitotic delay, also increased the number of polyploid cells, nuclear size, karyotype heterogeneity, chromosome bridge frequency, and micronucleus rates (Fig. 4G–4L). These results suggest that *YY2* overexpression might also increase CIN.

Given that BUB3 is a component of SAC, we overexpressed *BUB3* and examined its effect on CIN. *BUB3* overexpression induced cell death (Supplementary Fig. S7A) and increased CIN phenotypes (Supplementary Fig. S7B–S7E). These results conform with previous studies showing that SAC activation could also induce CIN and furthermore, cell death (12,34). To further confirm the causal relation between *YY2* overexpression-induced SAC hyperactivation and CIN, we suppressed SAC activity in HCT116 cells overexpressing *YY2*. Suppression of *BUB3* or treatment with reversine significantly decreased the number of polyploid cells (Supplementary Fig. S8A–S8B), chromosome bridge frequency (Supplementary Fig. S8C– S8D), and micronucleus rate (Supplementary Fig. S8E–S8F) in HCT116 cells overexpressing *YY2*, which is likely due to the suppression of SAC activity to near wild-type level in these cells as shown in Supplementary Fig. S5G-S5H. Together, our results clearly show that both *YY2* knockout and overexpression induced CIN.

### CIN levels determine tumor cell fates

The aforementioned results showing that both *YY2* knockout and overexpression induced CIN were intriguing, as *YY2* knockout suppressed SAC activity, shortened the mitotic time, and promoted HCT116 cell proliferation, whereas YY2 overexpression induced SAC hyperactivation, prolonged the mitotic time, enhanced cell death, and suppressed HCT116 cell proliferation. Meanwhile, clinical studies have shown that patients with high CIN have a better prognosis (5-7). Using TCGA CRC data set, we next performed correlation analysis between relapse-free survival rate of CRC patients treated with oxaliplatin, a drug clinically used for treating CRC, and the pretreatment levels of CIN70, a 70-gene signature that has been established as surrogate of CIN (35). The results showed that the moderate CIN levels, *i.e.*, the second and third quartiles, were associated with poor prognosis; meanwhile, high levels of CIN, *i.e.*, the fourth quartile, were associated with better survival, suggesting that excessive CIN might be deleterious for tumors (Fig. 5A).

**Figure 5.**
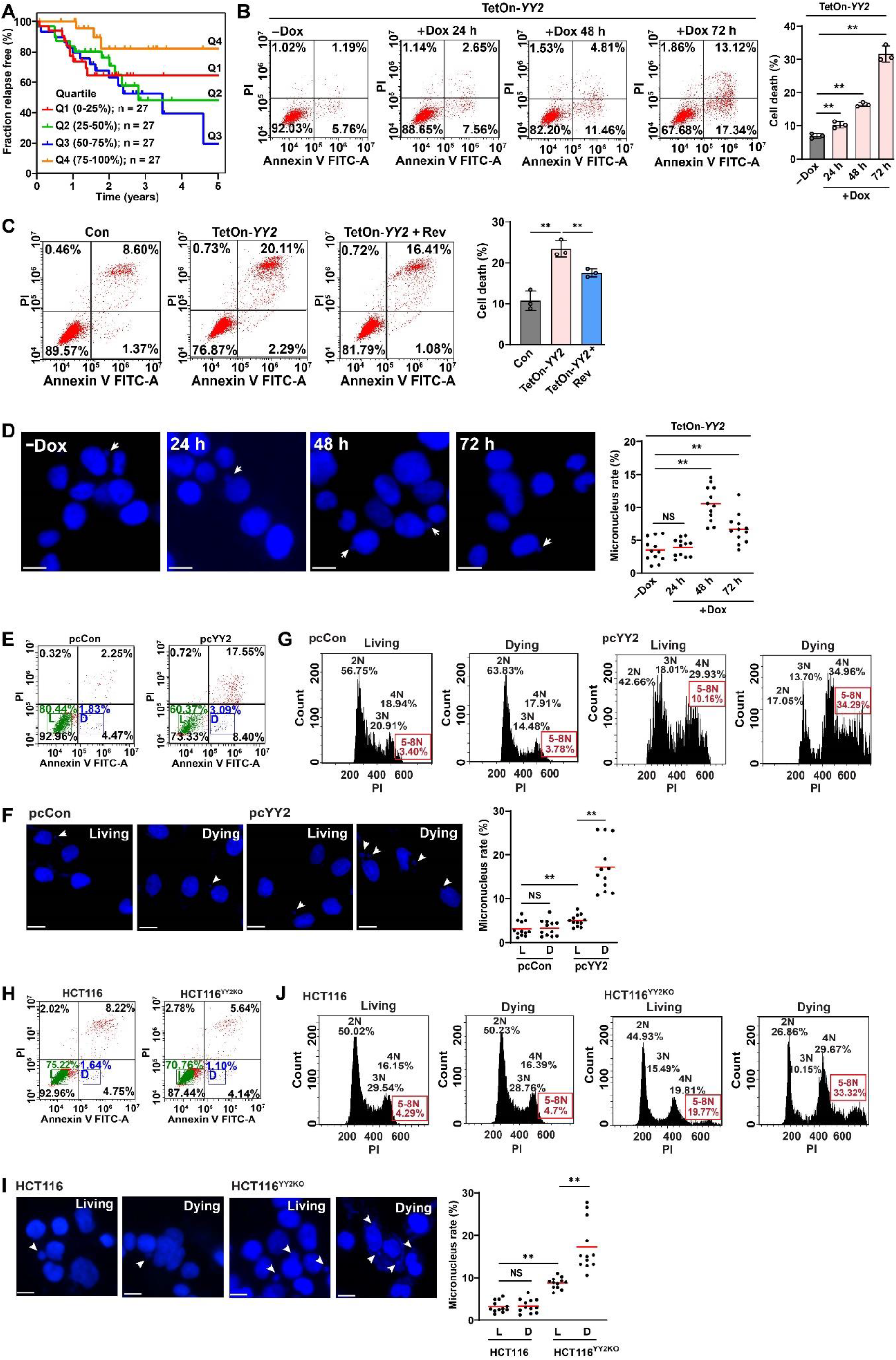
CIN level determines tumor cell fate. **A,** Kaplan-Meier relapse-free survival curves of 108 oxaliplatin-treated CRC patients stratified by CIN70 score quartile. **B–C,** Cell death rate of HCT116 cells after *YY2* induction by doxycycline (**B**) and after reversine treatment (final concentration: 0.2 μM) (**C**), as examined using Annexin V/PI staining. **D,** Micronucleus rate (scale bars: 20 μm; micronucleus rate: ratio of micronuclei number to total cell number; each dot represents micronucleus rate/slide with > 100 cells/slides; four technical replicates from three independent experiments) of HCT116 cells after *YY2* induction by doxycycline. **E–G,** Sorting (**E**), micronucleus rate (**F**), and DNA content (**G**) of living (L) and dying (D) *YY2*-overexpressed HCT116 cells. **H–J,** Sorting (**H**), micronucleus rate (**I**), and DNA content (**J**) of living (L) and dying (D) HCT116^YY2KO^ cells. Cells transfected with pcCon or wild-type HCT116 cells or infected with lentivirus generated using pTRIPZ-control (Con) were used as controls. Quantification data are shown as mean ± SD. All data were obtained from three independent experiments. *P* values were calculated by one-way ANOVA. Scale bars: 20 μm; pcCon: pcEF9-Puro; Dox: doxycycline (final concentration: 2 μg/ml); \*\**P* < 0.01; NS: not significant.

We next examined the fate of *YY2*-overexpressed cells. To control the timing of *YY2* overexpression, we established a Tet-On *YY2* overexpression system (Supplementary Fig. S9A–S9B). The cell death rate increased in a time-dependent manner after the induction of *YY2* overexpression (Fig. 5B); while treatment with reversine cancelled it (Fig. 5C), suggesting the correlation between *YY2* overexpression-induced SAC hyperactivation and cell death. Given that *YY2* overexpression also led to increased CIN (Fig. 4G–4L), these results indicated the possibility that YY2-induced cell death might be correlated with SAC hyperactivation-induced CIN. However, unlike the cell death rate, the micronucleus rate started to decline after reaching its peak at 48 h post-doxycycline treatment, leading to questions regarding the fate of cells with high CIN (Fig. 5D). To examine the relationship between the CIN level and cell death, we stained the cells with Annexin V and propidium iodide (PI), and sorted the “dying cells” (Annexin V^+^/PI^−^), in which nuclear and chromosomal DNA fragmentation had not occurred and the chromosome could still be observed, as well as “living cells” (Annexin V^-^/PI^−^) (Fig. 5E). Whereas there was no significant difference between the micronucleus rate of living and dying control cells, the micronucleus rate, as well as the number of cells with more than one micronucleus in dying *YY2*-overexpressed HCT116 cells, was significantly higher than that in living *YY2*-overexpressed cells (Fig. 5F). Similarly, whereas there was no significant difference in the percentage of polyploid cells between living and dying control cells (3.40% and 3.78%, respectively), the percentage of polyploid cells in dying *YY2*-overexpressed HCT116 cells was significantly higher than that in the corresponding living cells (34.29% and 10.16%, respectively; Fig. 5G).

Furthermore, while there were only few dying cells in non-treated HCT116^YY2KO^ cells, we confirmed that the micronucleus rate and percentage of polyploidy cells in those cells were significantly higher than in living HCT116^YY2KO^ cells (Fig. 5H-5J). Moreover, while knocking out *YY2* alone did not induce cell death and even benefits tumor cell proliferation, treatment with reversine, which could further weaken SAC activity as indicated by the shortened mitotic time (Supplementary Fig. S9C–S9D, Supplementary Video S10), increased CIN as well as cell death in HCT116^YY2KO^ cells (Supplementary Fig. S9E–S9F). Hence, these results further confirmed the association between cell death with high CIN. It is noteworthy that while excessive CIN eventually leads to cell death, intrinsic characteristics of the cells, such as different gene expression profiles, might also contribute to the capacity of the cells to tolerate CIN stress, and eventually to their different outcomes such as in cell proliferation (36,37). Indeed, we observed different expression profiles of genes related with DNA damage response and negative regulation of apoptosis in *YY2*-overexpressed and *YY2*-knocked out HCT116 cells (Supplementary Fig. S9G). Nevertheless, while further investigations are needed to systematically analyze these differences, our results clearly suggest that cells with excessive CIN induced by either YY2-mediated SAC hyperactivation or *YY2* knockout are more prone to death.

We further noticed that while the level of CIN in the living *YY2*-overexpressed HCT116 cells was significantly lower than in the dying *YY2*-overexpressed HCT116 cells, it is slightly higher than in living control cells, as indicated by the slightly increased micronucleus rate and percentage of polyploid cells (Fig. 5F–5G, respectively). As described above, moderate CIN level is correlated with poor prognosis (Fig. 5A). These results raised question regarding the characteristics of these moderate CIN cells. To examine these characteristics, we reversed *YY2* overexpression by withdrawing doxycycline after sorting the living cells to avoid continuous SAC hyperactivation (TetOn-*YY2* Trans; Supplementary Fig. S10A–S10B). CIN indicator analysis performed 2 days after doxycycline withdrawal showed that the slightly increased CIN level was maintained in living TetOn-*YY2* Trans cells (Fig. 6A–6E and Supplementary Fig. S10C), confirming that CIN is heritable and can be passed to the progeny cells as reported previously (38). Meanwhile, while continuous *YY2* overexpression suppressed the viability and colony-formation potential while increasing cell death rate (TetOn-*YY2* Conti), these characteristics recovered to the levels similar to those of control in TetOn-*YY2* Trans cells (Fig. 6F–6G and Supplementary Fig. S10D). However, under DNA-damage stress induced by oxaliplatin, TetOn-*YY2* Trans cells demonstrated robustly higher viability and colony-formation potential (Fig. 6H–6I). Moreover, the cell death rate of TetOn-*YY2* Trans cells was significantly lower than HCT116 cells, which were CIN-negative (Fig. 6J), suggesting that CIN level before oxaliplatin treatment contribute to the observed differences in viability between cells with continuous and transient *YY2* overexpression. Meanwhile, suppressing CIN in TetOn-*YY2* Conti cells using reversine decreased cell death and improved cell viability (Supplementary Fig. S10E–S10G), indicating that reducing excessive CIN level by inhibiting SAC in SAC-hyperactivated cells can enhance tumor cell drug resistance. Together, these data indicated the correlation between moderate CIN and tumor cell drug resistance.

**Figure 6.**
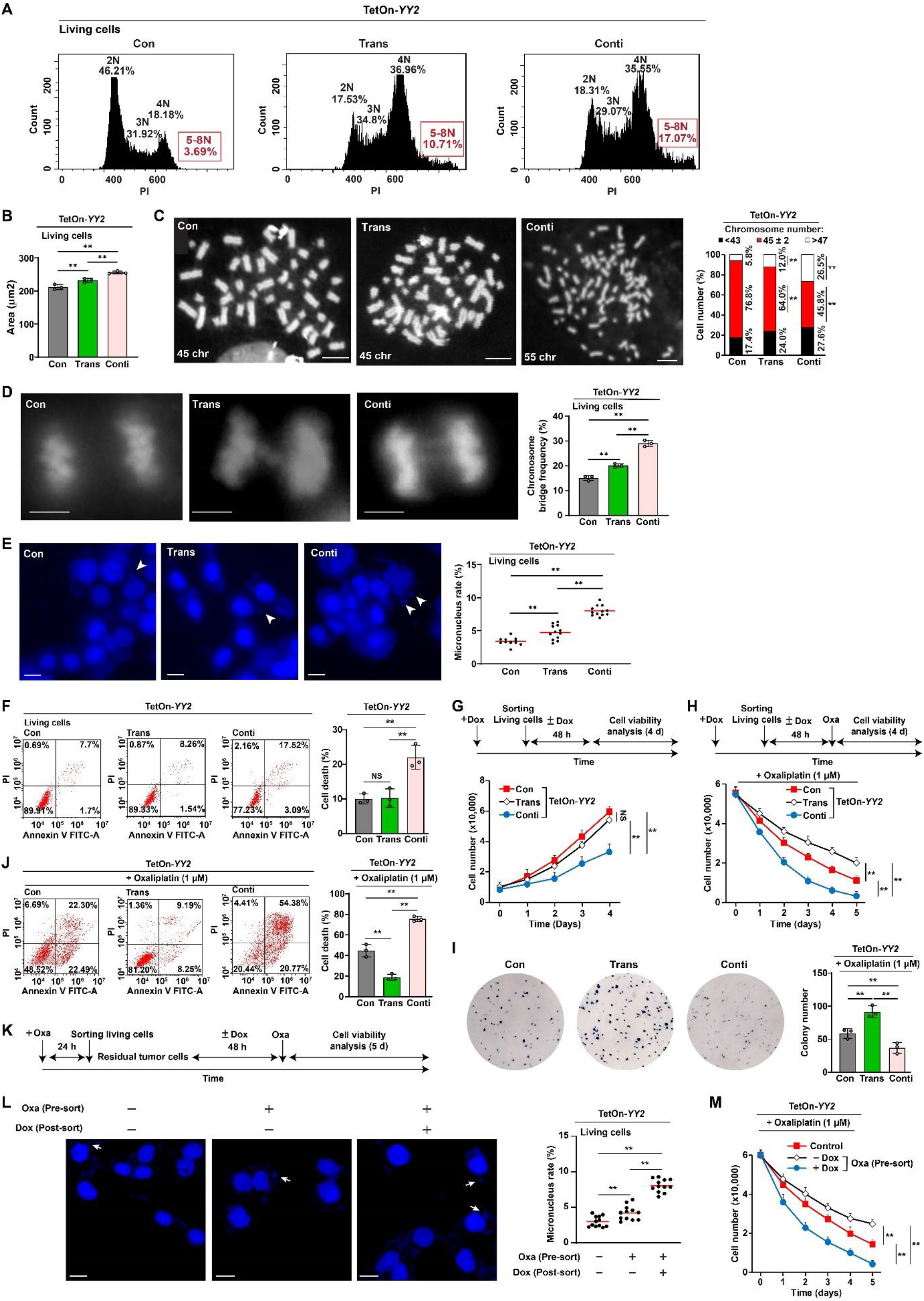
Residual tumor cells have moderate CIN and increased drug resistance. **A,** DNA content in HCT116 cells transiently overexpressing *YY2* (TetOn-*YY2* Trans cells) 48 h after sorting, as analyzed using PI staining and flow cytometry. **B,** Average nuclei size of TetOn-*YY2* Trans cells 48 h after sorting (each dot represents average nuclei size from one independent experiment, with total 100 cells/group). **C,** Chromosome number per cell in TetOn-*YY2* Trans cells 48 h after sorting, as analyzed using metaphase spread. Representative images (scale bars: 10 μm) and percentage of cells with indicated chromosome number (total cells counted: 50 cells/group, pooled from three independent experiments) are shown. **D,** Chromosome bridge frequency in TetOn-*YY2* Trans cells 48 h after sorting. Representative images (scale bars: 5 μm) and quantification results (each dot represents chromosome bridge frequency from one independent experiment, with total 100 mitotic-cells/group) are shown. **E,** Micronucleus rate of TetOn-*YY2* Trans cells 48 h after sorting. Representative images of micronuclei (indicated by arrowheads; scale bars: 20 μm) and micronucleus rate (ratio of micronuclei number to total cell number; each dot represents micronucleus rate/slide with > 100 cells/slides; four technical replicates from three independent experiments) are shown. **F–G,** Cell death rate at indicated time-points (**F**) and cell viability at 4 days after sorting (**G**) of TetOn-*YY2* Trans cells. **H–J,** Viability (**H**), colony-formation potential (**I**; each dot represents the mean value of three technical replicates), and cell death rate (**J**) of TetOn-*YY2* Trans cells at indicated times, 10 days, and 5 days after oxaliplatin treatment, respectively. **K–M,** Schematic diagram (**K**), micronucleus rate (**L**), and viability after oxaliplatin treatment (**M**) of residual HCT116 cells infected with TetOn-*YY2* lentivirus. Cells infected with control lentivirus generated from pTRIPZ-control (Con) were used as controls. TetOn-*YY2* Trans and Conti: cells infected with TetOn-*YY2* lentivirus and treated with doxycycline only before sorting or continuously, respectively. Pre-sort and post-sort: pre-sorting and post-sorting for living cells. Quantification data are shown as mean ± SD. All data were obtained from three independent experiments. *P* values were calculated by one-way ANOVA. Dox: doxycycline (final concentration: 2 μg/ml); \*\**P* < 0.01; NS: not significant.

Residual cells, which survived the anti-tumor drug treatment and undetectable by standard morphological examination during the remission phase between the cycles of chemotherapy, has attracted attention as one of the crucial factors of tumor drug resistance and tumor recurrence, and thus as a hurdle for anti-tumor therapy (39). The fact that living cells with YY2-induced SAC hyperactivation possessed moderate CIN suggests the possibility that moderate CIN is also involved in drug resistance in residual tumor cells that survived treatment with DNA damage-inducing agents. To test this possibility, we obtained residual tumor cells from CRC cells infected with TetOn-*YY2* lentivirus by sorting living cells that survived oxaliplatin treatment, treated them with or without doxycycline to induce *YY2* overexpression, and examined their viability under oxaliplatin treatment (Fig. 6K). Micronucleus assay revealed that without induction of *YY2* overexpression, residual cells that survived oxaliplatin treatment had moderate CIN level (Fig. 6L, middle row); furthermore, these cells demonstrated improved DNA-damage stress-resistance compared to that in control cells (Fig. 6M). Meanwhile, further induction of CIN by activating *YY2* overexpression in these residual cells induced excessive CIN (Fig. 6L, right row), leading to better drug sensitivity (Fig. 6M).

Our results demonstrated that YY2/SAC hyperactivation induces CIN in tumor cells, leading to the mortality of cells with excessive CIN. Moreover, our results suggest that moderate CIN, as observed in residual tumor cells surviving SAC hyperactivation or DNA damage-inducing agent treatment, is crucial for drug resistance in these cells; while increasing CIN in these cells could improve their drug sensitivity. Together, our findings highlight the crucial role of CIN levels in determining tumor cell fate and drug resistance.

### *YY2*-induced excessive CIN sensitizes tumor cells to DNA damage-inducing agent

Given that *YY2* overexpression induces excessive CIN, we examined its synergistic effect with a DNA damage-inducing agent. *YY2* overexpression significantly decreased the IC_50_ of oxaliplatin; in contrast, *YY2* knockout robustly increased it (Fig. 7A–7B). Furthermore, combinatorial index (CI) analysis using a previously described method (40,41) showed a synergistic effect between *YY2* overexpression and oxaliplatin treatment (Fig. 7C). The results of the EdU incorporation assay (Supplementary Fig. S11A–S11B) and cell death rate (Fig. 7D) showed that *YY2* overexpression further enhanced the suppressive effect of oxaliplatin on HCT116 cell proliferation potential while further promoting its effect on inducing cell death.

**Figure 7.**
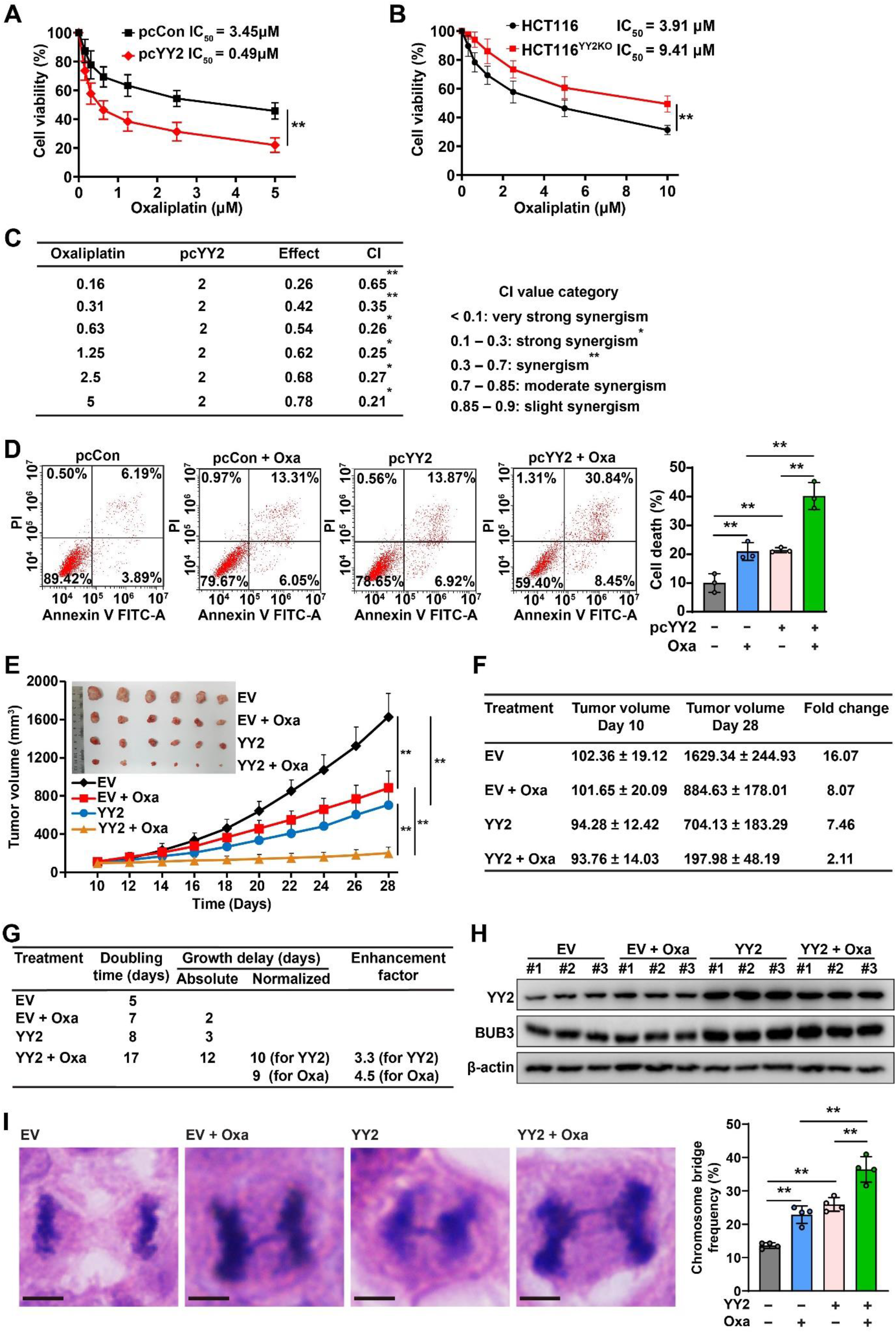
Excessive CIN sensitizes tumor cells to DNA-damage agents. **A–B,** Viability of *YY2*-overexpressed HCT116 cells (**A**) and HCT116^YY2KO^ cells (**B**) treated with indicated concentrations of oxaliplatin for 24 h. IC_50_ was calculated using CompuSyn. **C,** Combination index (CI) between oxaliplatin and *YY2* overexpression, as calculated using CompuSyn. **D,** Cell death rate of oxaliplatin-treated *YY2*-overexpressed HCT116 cells, as examined using Annexin V/PI staining and flow cytometry. **E,** Tumor volume and morphological images of xenografted tumors formed by *YY2*-overexpressed HCT116 cells and treated with oxaliplatin at indicated time-points (n = 6/group). **F,** Fold-change of xenografted tumor volumes at day 28 compared to those at the starting point of treatment (day 10). **G,** Enhancement factor of combinatory treatment of oxaliplatin and *YY2* overexpression. **H,** YY2 and BUB3 protein levels in xenografted tumors, as determined using western blotting. **I,** Chromosome bridge frequency in tissue sections of xenografted tumors. Representative images (scale bars: 5 μm) and quantification results (each dot represents chromosome bridge frequency from one slide, with total 100 mitotic-cells/group) are shown. β-actin was used as western blotting loading control. Cells transfected with pcCon, wild-type HCT116 cells, or cells infected with empty virus (EV) were used as controls. Quantification data are shown as mean ± SD. All data were obtained from three independent experiments. *P* values were calculated by one-way ANOVA. pcCon: pcEF9-Puro; \*\**P* < 0.01.

We then assessed the possibility of combining excessive CIN induced by *YY2* overexpression with an anti-tumor drug inducing DNA damage for anti-tumor therapeutic strategy. To this end, we established a *YY2*-overexpressed HCT116 stable cell line using a lentivirus and performed xenograft experiments (Supplementary Fig. S11C). Whereas the volume of xenografted tumors formed by control cells increased 16-fold within 4 weeks, *YY2* overexpression or oxaliplatin treatment alone suppressed this increase to approximately eight-fold. The suppressive effect was further enhanced by combining *YY2* overexpression and oxaliplatin treatment, which suppressed the increase in tumor volume to only two-fold (Fig. 7E–7F). The tumor morphology further confirmed this tendency (Fig. 7E). Furthermore, doubling time analysis showed that combining *YY2* overexpression and oxaliplatin treatment resulted in more than 9 days of growth delay compared to that with *YY2* overexpression or oxaliplatin treatment alone, thereby enhancing the therapeutic effect by approximately four-fold (Fig. 7G). Western blotting and immunohistochemistry (IHC) staining results showed that BUB3 expression was increased in tumors formed by *YY2*-overexpressed cells, whereas oxaliplatin did not affect YY2 and BUB3 expression (Fig. 7H and Supplementary Fig. S11D). Furthermore, IHC staining using γH2AX showed that DNA damage in tumor lesions, which was increased by *YY2* overexpression or oxaliplatin treatment alone, was further enhanced by combined oxaliplatin treatment and *YY2* overexpression (Supplementary Fig. S11D).

Finally, we examined CIN levels in the tumor lesions. Compared to that in the controls, *YY2* overexpression or oxaliplatin treatment increased the chromosome bridge frequency, whereas in oxaliplatin-treated *YY2*-overexpressed HCT116 cells, this was further increased (Fig. 7I). Collectively, these results demonstrated that *YY2* overexpression sensitizes tumor cells to oxaliplatin by increasing CIN, suggesting that this combination might be a potential anti-tumor therapeutic strategy. Together, our results demonstrated that YY2 is a novel SAC modulator that activates *BUB3* transcription, thereby influencing SAC activity. This subsequently determines tumor cell fate and drug resistance by regulating the levels of CIN (Fig. 8).

**Figure 8.**
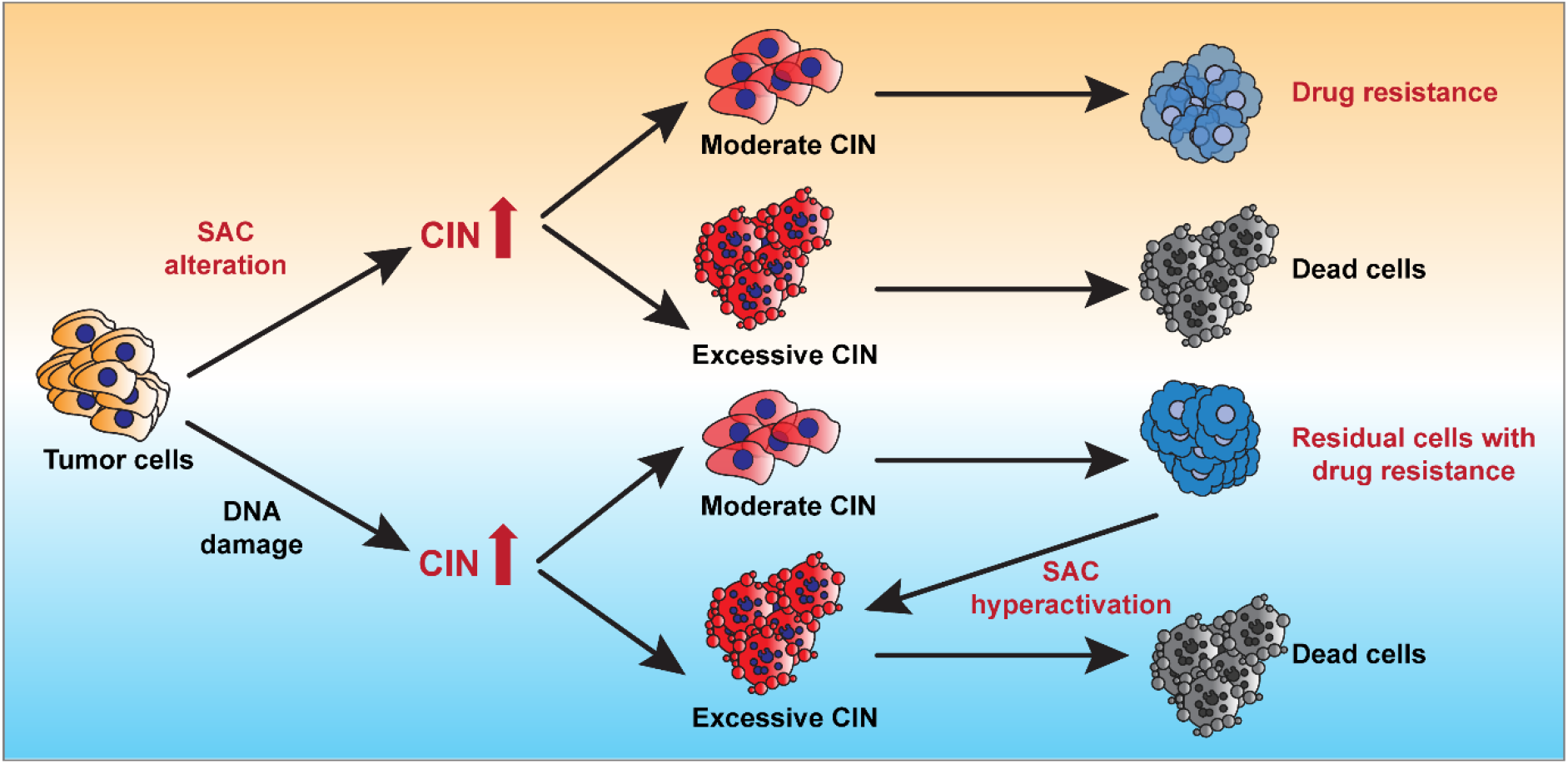
Schematic diagram showing the role of SAC activity and CIN level in determining tumor cell fate and drug resistance.

## Discussion

Impaired SAC is the main cause of chromosome segregation errors in mitosis, leading to numerical and structural chromosome changes in tumor cells (8,10,11). Mouse models have shown that impaired SAC promotes aneuploidy *in vivo* (42). Moreover, defective SAC activity is closely related to diseases such as tumors and mosaic variegated aneuploidy, a rare disorder with a high aneuploidy rate and increased tumor incidence (43). On the other hand, the role of SAC hyperactivation in tumorigenesis is still poorly understood, with conflicting reports on its pro- or anti-tumor effects (12,13,34). Nevertheless, mutations in SAC genes are rare in human tumors, indicating that they are not major contributors to the mitotic errors observed in tumor cells (14). In this study, we identified YY2, a transcription factor that oscillates during the cell cycle and peaks during the M phase, as a novel SAC modulator. YY2 directly activates the transcriptional activity of *BUB3*, a component of the SAC complex. We revealed that YY2 downregulation weakens SAC activity, thereby promoting tumorigenesis by accelerating sister chromatid segregation, mitotic exit, and tumor cell proliferation. This suggests a mechanism linking CIN to YY2 downregulation and loss-of-function mutations observed in various tumors (27,44). Meanwhile, *YY2* overexpression hyperactivates SAC, leading to prolonged mitotic time, decreased proliferation potential, and increased tumor cell death, thus suppressing tumorigenesis. YY2 downregulation in various tumors, including colorectal, prostate, ovarian, and liver cancers, is closely associated with poor survival and prognosis (44-46). While previous studies have revealed that YY2 can trigger ferroptosis, ultraviolet damage response, and p53-mediated cell cycle arrest (17,28,47), studies regarding its physiological and pathological functions are still very limited, and the mechanisms underlying its tumor suppressive effect have not been completely elucidated. Hence, our findings provide a new perspective regarding a novel function of YY2 in regulating CIN and subsequently tumor cell fate through its regulatory effect on SAC activity.

CIN is a hallmark of cancer and can drive tumorigenesis. Paradoxically, high CIN levels before treatment are associated with improved overall survival outcomes, whereas intermediate CIN levels are associated with poor clinical outcomes (5-7). A similar phenomenon was observed in bacteria and viruses, where higher genomic instability benefits the population in stressful environments through the development of mutations that provide a selective growth advantage; however, cells with drastic instability never become dominant in a population, as their instability levels lead to deleterious mutations exceeding the viability threshold (48). In tumor cells, this concept of a ‘just-right’ level of instability was also observed based on mathematical modeling of the evolutionary dynamics of genetically unstable populations (49,50), suggesting that whereas moderate CIN benefits tumorigenesis as it can allow tumor cells with varying genetic alterations to have greater chance of acquiring advantageous characteristics, such as increased proliferation, excessive CIN might induce tumor cell death. Yet, experimental evidence to support this hypothesis and the molecular mechanisms underlying tumor cell CIN are still lacking.

Intriguingly, *YY2* overexpression, which hyperactivates the SAC and triggers tumor suppressive effects, also induces CIN. Indeed, previous studies have revealed that SAC hyperactivation, for example by overexpression of another SAC component, *MAD2*, or by knocking down *p31*, a component of SAC silencing, could also induce CIN and cell death (13,34,51,52). Here we revealed that in the *YY2*-induced SAC-hyperactivated cell population, dying cells had significantly higher CIN than living cells, thus provides evidence that high CIN induces cell death and is thus deleterious for tumor cells. Meanwhile, although significantly lower than that in dying cells, living cells from this population also exhibited a certain degree of CIN and improved DNA-damage resistance compared to those in control cells, suggesting that moderate CIN is beneficial for tumorigenesis. Together with the fact that knocking out *YY2* alone is not sufficient for inducing cell death, while further inhibiting SAC activity in *YY2*-knocked out cells could also lead to excessive CIN and cell death, our findings clearly showed that different levels of CIN is crucial for determining tumor cell fate. Obviously, while excessive CIN will eventually lead to cell death, intrinsic characteristics of the cells, such as their gene expression profiles in responses to DNA damage stress and apoptotic stimuli, also contribute greatly to their capacity in tolerating CIN stress, and subsequently, the threshold of CIN-induced cell death. Hence, while further systematical investigation is needed to reveal the way tumor cells explore their fitness landscape upon CIN induction, our findings provide experimental evidence that conforms to the ‘just-right’ model.

DNA damage-inducing agents such as oxaliplatin, taxol, and radiotherapy, which have been used clinically for treating tumors, can induce mitotic errors and CIN (6,53); while SAC activator, such as MK-1775 and ZN-c3, are in clinical trials and are considered as potential anti-tumor drugs (54). Meanwhile, increasing evidences suggest that a population of tumor cells remain viable after exposure to various anti-tumor drugs. These residual cells display reduced drug sensitivity, thereby providing a reservoir of cells that might seed the growth of drug-resistant recurrent tumor. Therefore, targeting residual cells have attracted attention as a crucial point for anti-tumor therapeutic strategy; however, the mechanisms by which these cells are generated are still poorly understood. Our findings showed that residual tumor cells, that survived DNA damage-inducing agents as well as YY2-induced SAC hyperactivation, possess moderate CIN and are more resistant. Hence, while the underlying mechanism needs to be further elucidated, our results link up for the first time the level of CIN and residual tumor cells as described by the ‘just-right’ model (55,56), thus providing new perspective regarding the generation of residual tumor cells as well as their drug resistance. Furthermore, although further study is needed, these findings also point to possible complications, such as predisposing normal or benign tumor cells to becoming more tumorigenic, owing to the induction of moderate CIN mediated by DNA damage-based anti-tumor therapies. Moreover, these findings indicate the possibility of applying the pretreatment CIN level as an indicator to identify patients who would benefit from DNA damage-based anti-tumor therapies.

Subsequently, our results also demonstrated that *YY2* overexpression-induced SAC hyperactivation, which triggers excessive CIN, significantly sensitizes CRC cells to oxaliplatin. Combining *YY2* overexpression and oxaliplatin treatment significantly delayed tumorigenesis and enhanced the therapeutic effect in a xenograft mouse model, most plausibly by increasing CIN, as results showed synergism between YY2/SAC hyperactivation-induced excessive CIN and DNA damage-inducing agents. Therefore, our findings suggest a potential anti-tumor combinatory therapeutic strategy based on a DNA damage-inducing agent and YY2-induced excessive CIN, to improve the efficacy of the former, and at the same time, preventing the generation of residual tumor cells surviving DNA damage-inducing agents with moderate CIN, which could trigger drug resistance.

Taken together, we identified YY2 as a novel modulator of SAC activity in which alterations could induce different degrees of CIN and determine tumor cell fates. This provides experimental evidence explaining the CIN paradox in cancer. Furthermore, our findings have unraveled a mechanism of drug resistance in residual cells surviving anti-tumor therapies such as those using DNA damage-inducing agents and SAC activation, and suggest a novel anti-tumor therapeutic strategy that combines SAC activity regulators and DNA damage-inducing agents.

## Data availability

All raw data generated in this study are available upon request from the corresponding author. RNA-Seq data of *YY2*-overexpressed cells used here was obtained from our previous study. The data was previously deposited in Gene Expression Omnibus and are available at GSE184138. ChIP-Seq data analyzed in this study were obtained from Gene Expression Omnibus at GSE76856. TEX14 tissues expression was obtained from Genotype-Tissue Expression project at https://gtexportal.org/home/gene/TEX14. TCGA CRC data set was obtained from cBioportal (coadread_tcga_pan_can_atlas_2018).

## Acknowledgments

The authors would like to thank Professor Xia Zhang (Institute of Pathology, Southwest Hospital, Third Military Medical University) for his helpful comments in preparing this manuscript.

## Author contributions

V.K. and S.W. conceptualized and supervised this study; R.H., I.T.S.M., W.D., and W.L. performed the experiments; R.H., I.T.S.M., W.D., M.W., S.H., M.M., V.K., and S.W. analyzed and interpreted the data; R.H., W.D., V.K., and S.W. wrote the paper; H.Z. collected human clinical samples and performed clinical samples analysis; M.M. predicted and constructed shRNA expression vectors.

