## Supplemental Files for "YY2/BUB3 axis-mediated SAC hyperactivity determines tumor cell fate through chromosomal instability"

**This PDF file includes:**

Supplementary Materials and Methods.

Figure S1. YY2 regulates M phase progression.

Figure S2. YY2 induces degradation of cyclin B and securing proteins.

Figure S3. SAC is crucial for YY2 tumor suppressor activity.

Figure S4. YY2 enhances BUB3 mRNA expression levels.

Figure S5. YY2 regulates mitosis and tumorigenic potential through BUB3-induced SAC hyperactivation.

Figure S6. Alteration in YY2 expression induces CIN.

Figure S7. *BUB3* overexpression induces CIN.

Figure S8. *YY2* overexpression induces CIN by hyperactivating SAC.

Figure S9. SAC inhibition induces excessive CIN in *YY2*-knockout cells.

Figure S10. *YY2*-transient overexpression induces heritable CIN.

Figure S11. YY2 enhances oxaliplatin tumor suppressive activity.

Figure S12. Uncropped western blots with the indicated areas of selection in Figs. 1, 3, 7, and Supplementary Figs. S1, S2, S3, S4, S5, S9, S10, S11.

Table S1. Primer pairs used for qRT-PCR.

Table S2. Antibodies used for western blotting, immunohistochemistry, and ChIP assay.

Video Legends S1-S10

**Other Supplementary Materials for this manuscript include the following:**

Videos S1 to S10

**Supplementary Materials and Methods**

**RNA sequencing and data analysis**

RNA-seq analysis were performed by Shanghai Bio Technology Corporation (Shanghai, China) using Illumina HiSeq 2500 (Illumina, San Diego, CA; three replicates for each group). Sequencing raw reads were pre-processed by filtering out rRNA reads, sequencing adapters, short-fragment reads and other low-quality reads. Tophat v2.1.0 was used to map the cleaned reads to human reference genome ensemble GRCh38 (hg38) with two mismatches. After genome mapping, Cufflinks v2.1.1 was run with a reference annotation to generate FPKM values for known gene models. Differentially expressed genes were identified using Cuffdiff. The *P*-value significance threshold in multiple tests was set by the false discovery rate (FDR). The fold-changes were also estimated according to the FPKM in each sample. To identify DEGs in *YY2*-overexpressed cells, pairwise comparisons between each transfected sample and each control sample were first performed to identify genes upregulated or downregulated in each *YY2*-overexpressed sample with an FDR $\leq 0.05$ relative to each control.

**Tumor growth delay and enhancement factor**

Tumor growth delay and enhancement factor was calculated using the following equation: (1): absolute growth delay was calculated by subtracting the doubling time of the tumor in the treated group from that of the EV group; (2): normalized growth delay for YY2 group was calculated by subtracting the absolute growth delay of the YY2 + oxaliplatin group from that of the EV + oxaliplatin group; (3): normalized growth delay for oxaliplatin was calculated by subtracting the absolute growth delay of the YY2 + oxaliplatin group from that of the YY2 group; (4): enhancement factor for YY2 group was calculated by dividing the normalized growth delay for

YY2 group (2) with the absolute growth delay of the YY2 group; (5): enhancement factor for oxaliplatin was calculated by dividing the normalized growth delay for oxaliplatin (3) with the absolute growth delay of the EV + oxaliplatin group.

**CIN70 scoring and survival data analysis**

CIN70 scores were calculated as the mean expression of the 70 probe sets matching the 70 genes of the CIN70 signature. The expression level of the 70 genes of CIN70 was obtained from TCGA dataset of CRC patients treated with oxaliplatin (cBioportal; coadread_tcga_pan_can_atlas_2018). All CRC samples were stratified into CIN70 quartiles according to the CIN70 scores.

**Cell sorting**

Cells were stained with Annexin V/PI according to manufacturer's instruction (Neobiosciences, Shanghai, China), and sorted with MoFlo XDP cell sorter (Beckman Coulter, Indianapolis, IN) for living (Annexin $V^-/PI^-$) and dying ($Annexin^+/PI^-$) cells. Data was analyzed using CytExpert (Beckman Coulter).

**Preparation of cells transiently overexpressing *YY2***

*YY2*-inducible cell line were treated with doxycycline (final concentration: 2 μg/ml) for at 48 h prior to cell sorting. Following cell sorting, the collected cells were cultured in medium without doxycycline for 48 h to turn off *YY2* overexpression.

**Cell death assay**

Cells were prepared as described above and re-seeded in a 6-well plate ($3 \times 10^5$ cells/well). Cells were stained with Annexin V/PI (NeoBiosciences) according to manufacturer's instruction 24 h after transfection and subjected to flow cytometry. Data was analyzed using CytExpert software.

**Cell viability assay and calculation of $IC_{50}$**

Cells were prepared as described above, re-seeded into 96-well plates ($5 \times 10^3$ cells/well), and treated with oxaliplatin (final concentration: 1 μM). Cell numbers were measured by colorimetric assay with MTS (Promega) at the indicated time points. $IC_{50}$ was calculated based on the results of cell viability at different dose of drugs (final concentrations: 0.16 μM, 0.31 μM, 0.63 μM, 1.25 μM, 2.5 μM, 5 μM, and 10 μM) using Compusyn® (www.combosyn.com; Combosyn Inc. Paramus, NJ). CI values were calculated using Compusyn® based on corresponding cell viability assay results using the Chou-Talalay methods. CI values were categorized as follow: < 0.1: very strong synergism; 0.10–0.30: strong synergism; 0.30–0.70: synergism; 0.70–0.85: moderate synergism; 0.85–0.90: slight synergism; 0.90–1.10: nearly additive; 1.10–1.20: slight antagonism; 1.20–1.45: moderate antagonism; 1.45–3.30: antagonism; 3.30–10: strong antagonism; > 10: very strong antagonism.

**Preparation and analysis of residual tumor cells**

*YY2*-inducible cell line was prepared as described above and cultured without doxycycline induction. Cells were treated with oxaliplatin (final concentration: 1 μM) for 24 h prior to sorting for living cells as described above. Oxaliplatin was withdrawn after cell sorting, and living cells were further cultured in medium with or without doxycycline (final concentration: 2 μg/ml) for 48

h. Cells were treated with oxaliplatin (final concentration: 1 μM) for 5 days, and cell viability was analyzed as described previously.

For micronucleus analysis, living cells were further cultured with or without doxycycline (final concentration: 2 μg/ml) for 48 h after sorting. Micronucleus staining was performed as described above.

**5-ethynyl-2′-deoxyuridine (EdU) incorporation assay and colony-formation assay**

Cells were prepared as described above and then re-seeded in 48-well plate ($5 \times 10^4$ cells/well). EdU incorporation and staining were performed using BeyoClick™ EdU Cell Proliferation Kit with Alexa Fluor 488 (Beyotime Biotechnology) according to the manufacturer's instruction. Nuclei were stained with Hoechst. Images were taken with fluorescence microscope (Olympus IX71). Quantification of EdU-positive and Hoechst-positive cells was performed using ImageJ and the results are shown as the ratio of EdU-positive cells to Hoechst-positive cells.

For colony-formation assay, 300 cells were cultured in a 6-well plate for 8 days. Cells were then fixed with 4% paraformaldehyde and stained with methylene blue. The colonies were then counted. The investigator was blinded during the assessment.

**Dual luciferase reporter assay**

Cells were seeded into 24-well plates ($8 \times 10^4$ cells/well). Twenty-four hours later cells were co-transfected with the indicated overexpression vector, reporter vector, and *Renilla* luciferase expression vector (pRL-SV40, Promega) as the internal control. Luciferase activities were measured using Dual Luciferase Assay System (Promega) 48 h after transfection. Firefly luciferase activities were normalized with the corresponding *Renilla* luciferase activities.

**Chromatin immunoprecipitation (ChIP) assay**

Chromatin was immunoprecipitated using ChIP Assay Kit (Beyotime Biotechnology) according to the manufacturer′s instructions. Briefly, cells were lysed, and chromatin was immunoprecipitated using protein A + G agarose/salmon sperm DNA and anti-YY2 antibody or normal rabbit IgG, uncrosslinked for 4 h at 65 °C, and treated with 0.5 M EDTA, 1 M Tris (pH 6.5) and 20 mg/ml proteinase K. Immunoprecipitated chromatin was then subjected to PCR with PrimeSTAR Max (Takara Bio). Primer sequences for amplifying *BUB3* promoter region containing the predicted YY2-binding site were 5′-GCTGTCGTTTCAGGACCCTT-3′ (forward primer) and 5′-AATCAGACCAGCCTTTGCCC-3′ (reverse primer).

**RNA extraction and quantitative real-time PCR (qRT-PCR)**

Total RNA was extracted using TRIzol (Invitrogen Life Technology) according to the manufacturer's instructions. Total RNA (1 μg) was then reverse transcribed into cDNA using a PrimeScript Reagent Kit with gDNA Eraser (Takara Bio). qRT-PCR was performed using SYBR Premix Ex Taq (Takara Bio). The sequences of the primers used are listed in Table S1. β-actin was used to normalize sample amplifications.

For absolute qRT-PCR, purified PCR amplicons with concentrations ranging from $1 \times 10^1$ to $1 \times 10^7$ copies/ml were used as standards. Briefly, amplicons were purified by gel electrophoresis followed by purification using Universal DNA Purification Kit (Tiangen Biotech). The concentration of the purified amplicons were measured using NanoDrop (Thermo Scientific, Waltham, MA). Concentration of the DNA copies for each amplicons were obtained using the following equation: DNA (copies/μl) = 6.02 x $10^{23}$ (copies/mol) × DNA concentration (g/μl) / [DNA length (bp) × 660 (g/mol/bp)].

##### Western blotting

Total cells were lysed with RIPA lysis buffer supplemented with a protease inhibitor and phosphatase inhibitor cocktail (complete cocktail, Roche Applied Science, Mannheim, Germany). For samples from xenografted tumors, frozen specimens were homogenized with RIPA lysis buffer with protease inhibitor and phosphatase inhibitor cocktail to obtain protein extracts. Samples with equal amounts protein were electrophoresed on sodium dodecyl sulfate-polyacrylamide gels before being transferred to a polyvinylidene fluoride membrane with 0.45-μm pore size (Millipore, Billerica, MA). Antibodies used are listed in Table S2, and immunoblotting with an anti-β-actin antibody was conducted to ensure equal protein loading. Signals were detected using the SuperSignal West Femto Maximum Sensitivity Substrate detection system (Thermo Scientific).

##### Protein degradation assay

For protein degradation assay, cells were synchronized at prometaphase by nocodazole treatment, as described above. Mitotic cells were washed two times in PBS and either collected immediately (0 h) or released in drug-free medium. Protein samples were collected at the indicated time points and were subjected to western blotting as described above. Protein's half-life was determined by quantifying western blotting results using ImageJ.

##### Immunohistochemistry and Hematoxylin-Eosin staining

Fresh xenografted tumor lesions were fixed using 4% paraformaldehyde for overnight, embedded in paraffin, and sectioned at 4 μm thickness using a cryostat. After being dewaxed using xylene and rehydrated, sections were subjected to immunohistochemical staining. Briefly, the tissue sections were incubated with primary antibodies for 1 h. The specimens were then incubated with corresponding second antibodies conjugated with horse-radish peroxidase. Visualization was performed using a DAB Kit (DAKO, Beijing, China) under microscope. The nuclei were then counterstained with hematoxylin (Beyotime Biotechnology), then the sections were dehydrated and mounted with coverslip. Antibodies used were listed in Supplementary Table S2. Images were taken by using Pannoramic Midi (3DHistech, Budapest, Hungary).

#### Supplementary Figure 1

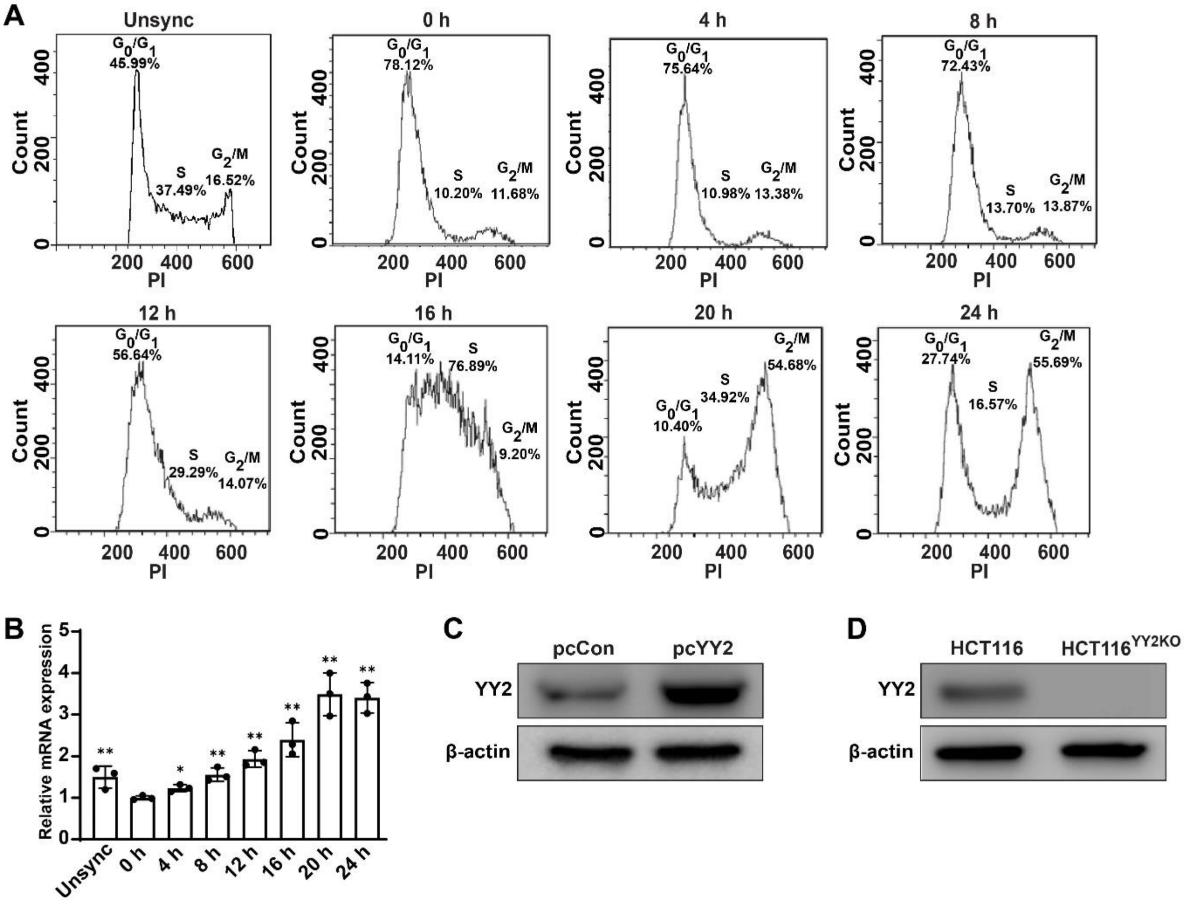


**Figure S1. YY2 regulates M phase progression.** (**A**) Analysis of cell cycle progression at indicated time-points starting immediately after serum starvation release, as examined using PI staining and flow cytometry. Representative images are shown. (**B**) YY2 mRNA expression level in HCT116 cells at indicated time-points after serum starvation release, as analyzed using qRT-PCR. YY2 mRNA levels in each time-point were shown as relative to that at 0 h after serum starvation release, which was assumed as 1. (**C**–**D**) YY2 protein expression level in HCT116 cells transfected with *YY2* overexpression vector (C) and in HCT116$^{YY2KO}$ cells (D), as determined using western blotting. Cells transfected with pcCon or wild-type HCT116 cells were used as controls. β-actin was used for qRT-PCR normalization and as western blotting loading control. Quantification data are shown as mean ± SD. All data were obtained from three independent experiments. *P* values were calculated by one-way ANOVA. Unsync: unsynchronized cells; pcCon: pcEF9-Puro; * $P < 0.05$; ** $P < 0.01$.

#### Supplementary Figure 2

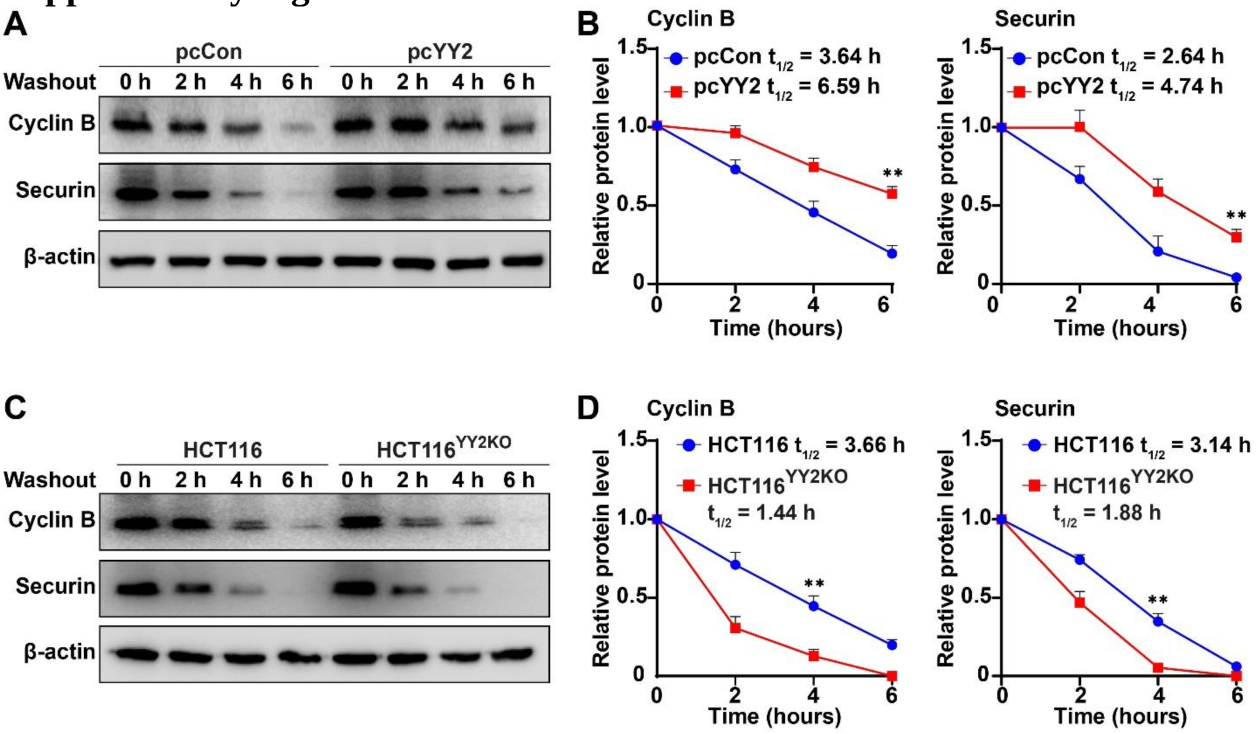


**Figure S2. YY2 induces degradation of cyclin B and securin. (A–B)** Cyclin B and securin protein expression levels in *YY2*-overexpressed HCT116 cells at indicated time-points after nocodazole washout, as analyzed using western blotting. Representative images (A) and the quantification results (B) are shown. **(C–D)** Cyclin B and securin protein expression levels in $HCT116^{YY2KO}$ cells at indicated time-points after nocodazole washout, as analyzed using western blotting. Representative images (C) as well as quantification results (D) are shown. Cells transfected with pcCon or wild-type HCT116 cells were used as controls. β-actin was used for western blotting loading control. Quantification data are shown as mean ± SD. All data were obtained from three independent experiments. *P* values were calculated by one-way ANOVA. pcCon: pcEF9-Puro; $^{**}P < 0.01$.

#### Supplementary Figure 3

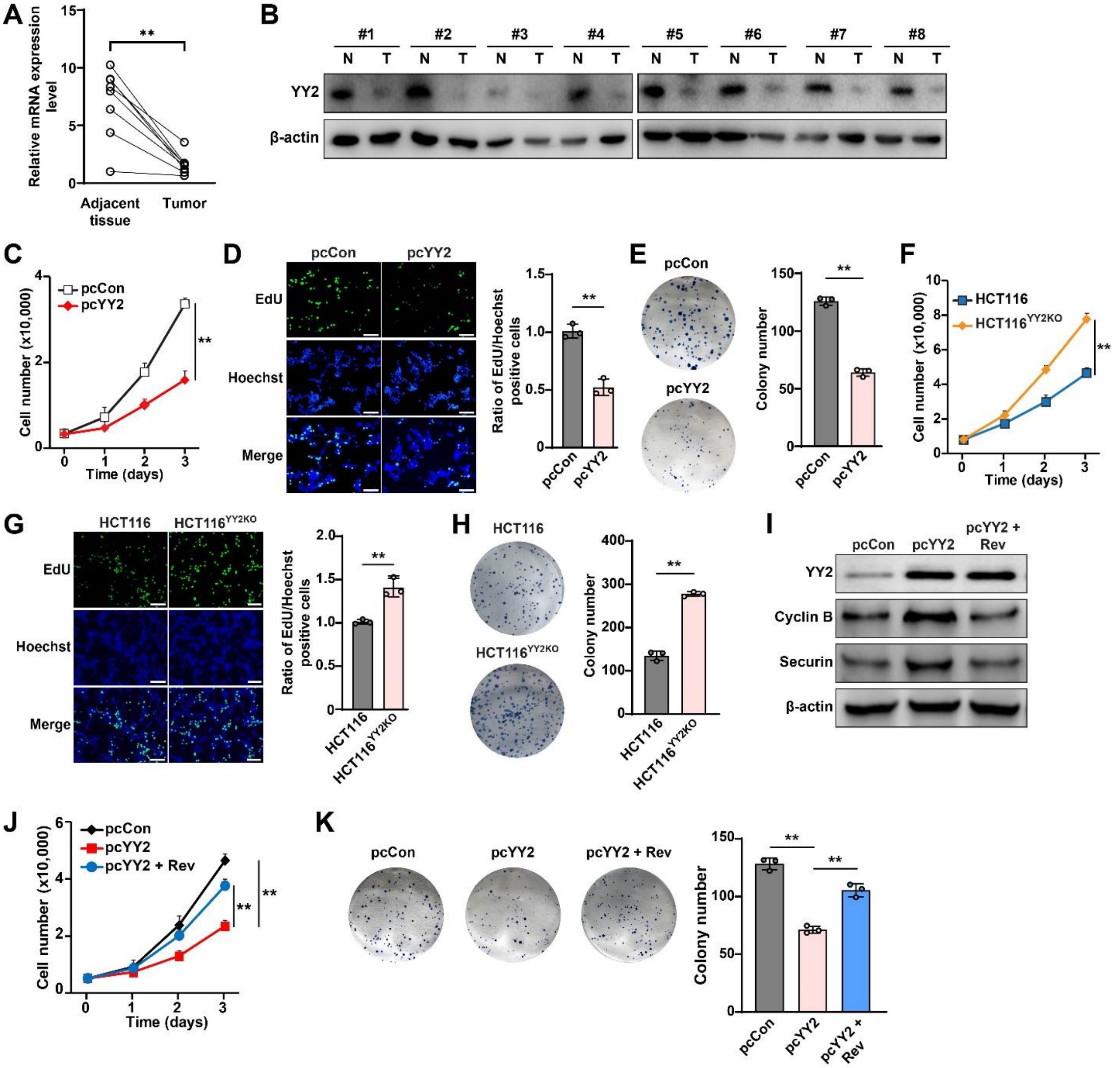


**Figure S3. SAC is crucial for YY2 tumor suppressor activity.** (**A–B**) YY2 mRNA (A; n = 16) and protein (B; n = 8) expression levels in clinical human colorectal cancer and corresponding normal adjacent tissues, as analyzed using qRT-PCR and western blotting, respectively. (**C**) Viability of *YY2*-overexpressed HCT116 cells at indicated time-points. (**D**) Proliferation potential of *YY2*-overexpressed HCT116 cells, as determined using EdU-incorporation assay. Representative images (left; scale bars: 200 μm) and ratio of proliferative cells (right; each dot represents the mean value of three technical replicates) are shown. (**E**) Colony-formation potential of *YY2*-overexpressed HCT116 cells. Representative images (left) and colony numbers (right; each dot represents the mean value of three technical replicates) are shown. (**F**) Viability of HCT116$^{YY2KO}$ cells at indicated time-points. (**G–H**) Proliferation (G) and colony-formation potentials (H) of HCT116$^{YY2KO}$ cells. (**I**) Cyclin B and securin protein expression levels in HCT116 cells overexpressing *YY2* and treated with reversine, as determined using western blotting. (**J**) Viability of HCT116 cells overexpressing *YY2* and treated with reversine at indicated time points. (**K**) Colony-formation potential of HCT116 cells overexpressing *YY2* and treated with reversine. Cells transfected with pcCon or wild-type HCT116 cells were used as controls. β-actin was used as western blotting loading control. Quantification data are shown as mean ± SD. All data were obtained from three independent experiments. *P* values were calculated by one-way ANOVA. pcCon: pcEF9-Puro; Rev: reversine (final concentration: 0.2 μM); $^{**}P < 0.01$.

#### Supplementary Figure 4

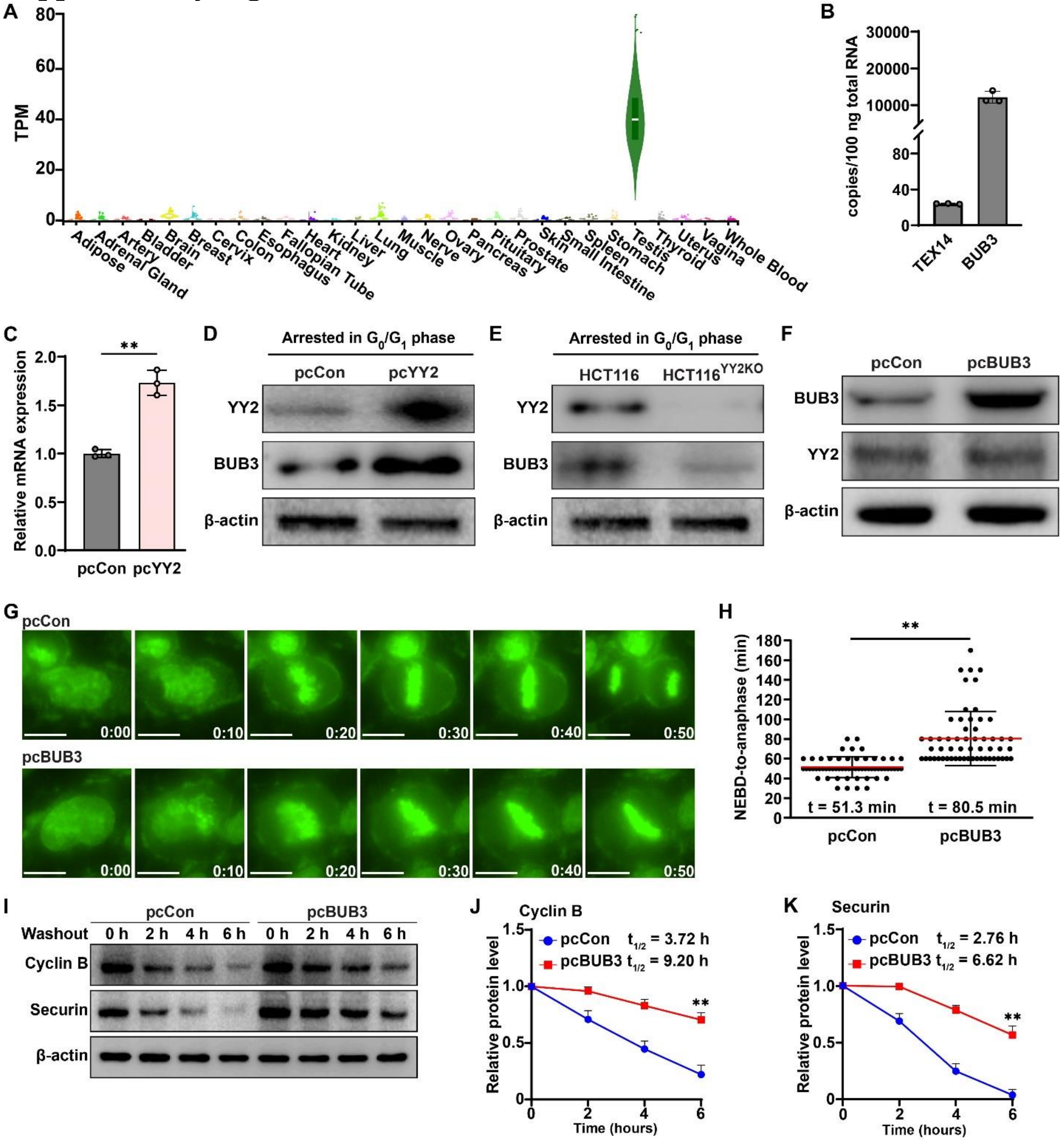


**Figure S4. YY2 enhances *BUB3* mRNA expression levels**. (**A**) GTEx tissue-wide expression profiles of TEX14. (**B**) Copy numbers of TEX14 and BUB3 mRNA in HCT116 cells, as analyzed using absolute qRT-PCR. (**C**) BUB3 mRNA expression level in HCT116 cells overexpressing *YY2*, as analyzed using qRT-PCR. (**D–E**) BUB3 protein expression level in *YY2*-overexpressed (D) and *YY2*-knockout (E) HCT116 cells arrested at $G_0/G_1$ phase following continuous serum starvation treatment, as determined using western blotting. (**F**) BUB3 protein expression level in HCT116 cells transfected with *BUB3* overexpression vector, as determined using western blotting. (**G–H**) Mitotic time of *BUB3*-overexpressed HCT116 cells, as determined using time-lapse microscopy. Representative images (G; scale bars: 20 μm) and scatter plot showing the time-length from NEBD to anaphase (H; n = 60, pooled from three independent experiments). (**I**–**K**) Cyclin B and securin protein expression levels in *BUB3*-overexpressed HCT116 cells at indicated time-points after nocodazole washout, as analyzed using western blotting. Representative images (I) as well as quantification results of cyclin B (J) and securin (K) are shown. Cells transfected with pcCon or wild-type HCT116 cells were used as control. β-actin was used for qRT-PCR normalization and as western blotting loading control. Quantification data are shown as mean ± SD. All data were obtained from three independent experiments. *P* values were calculated by one-way ANOVA. pcCon: pcEF9-Puro; $^{**}P < 0.01$.

#### Supplementary Figure 5

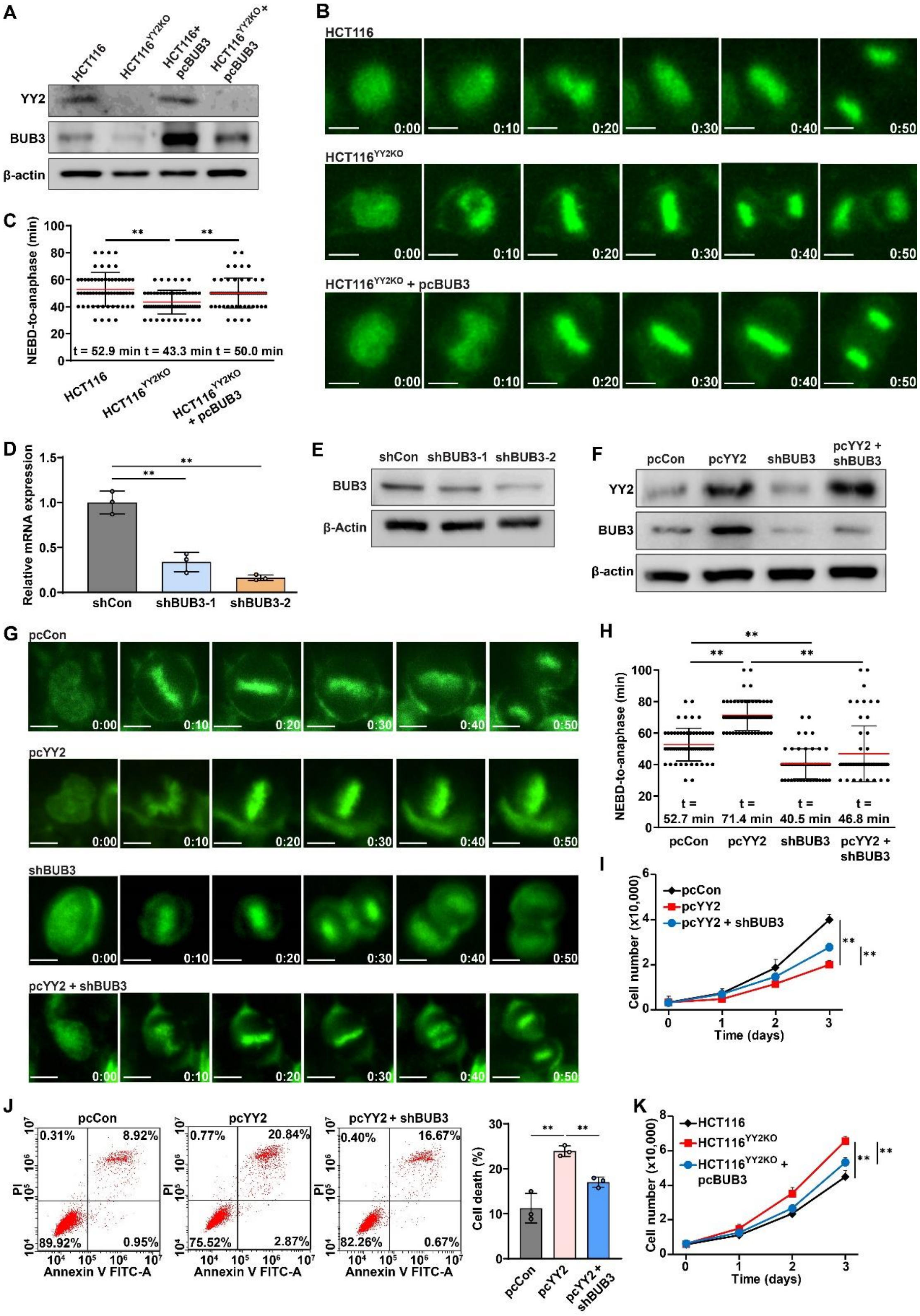

**Figure S5. YY2 regulates mitosis and tumorigenic potential through BUB3-induced SAC hyperactivation.** (**A**) BUB3 and YY2 protein expression levels in *BUB3*-overexpressed

HCT116$^{YY2KO}$ cells, as determined using western blotting. (**B–C**) Mitotic time of *BUB3*-overexpressed HCT116$^{YY2KO}$ cells, as determined using time-lapse microscopy. Representative images (B; scale bars: 20 μm) and scatter plot showing the time-length from NEBD to anaphase (C; n = 60, pooled from three independent experiments). (**D–E**) BUB3 mRNA (D) and protein (E) expression levels in HCT116 cells transfected with shRNA expression vectors targeting different sites of *BUB3*, as analyzed using qRT-PCR and western blotting, respectively. (**F**) BUB3 and YY2 protein expression levels in *BUB3* knocked-down, *YY2*-overexpressed HCT116 cells, as determined using western blotting. (**G–H**) Mitotic time of *BUB3* knocked-down, *YY2*-overexpressed HCT116 cells, as determined using time-lapse microscopy. Representative images (G; scale bars: 20 μm) and scatter plot showing the time-length from NEBD to anaphase (H; n = 60, pooled from three independent experiments). (**I–J**) Viability (I) and cell death rate (J) of *BUB3* knocked-down, *YY2*-overexpressed HCT116 cells at indicated time points. (**K**) Viability of *BUB3*-overexpressed HCT116$^{YY2KO}$ cells at indicated time points. Cells transfected with pcCon, shCon, or both pcCon and shCon were used as controls. β-actin was used for qRT-PCR normalization and as western blotting loading control. Quantification data are shown as mean ± SD. All data were obtained from three independent experiments. *P* values were calculated by one-way ANOVA. pcCon: pcEF9-Puro. $^{**}P < 0.01$.

#### Supplementary Figure 6

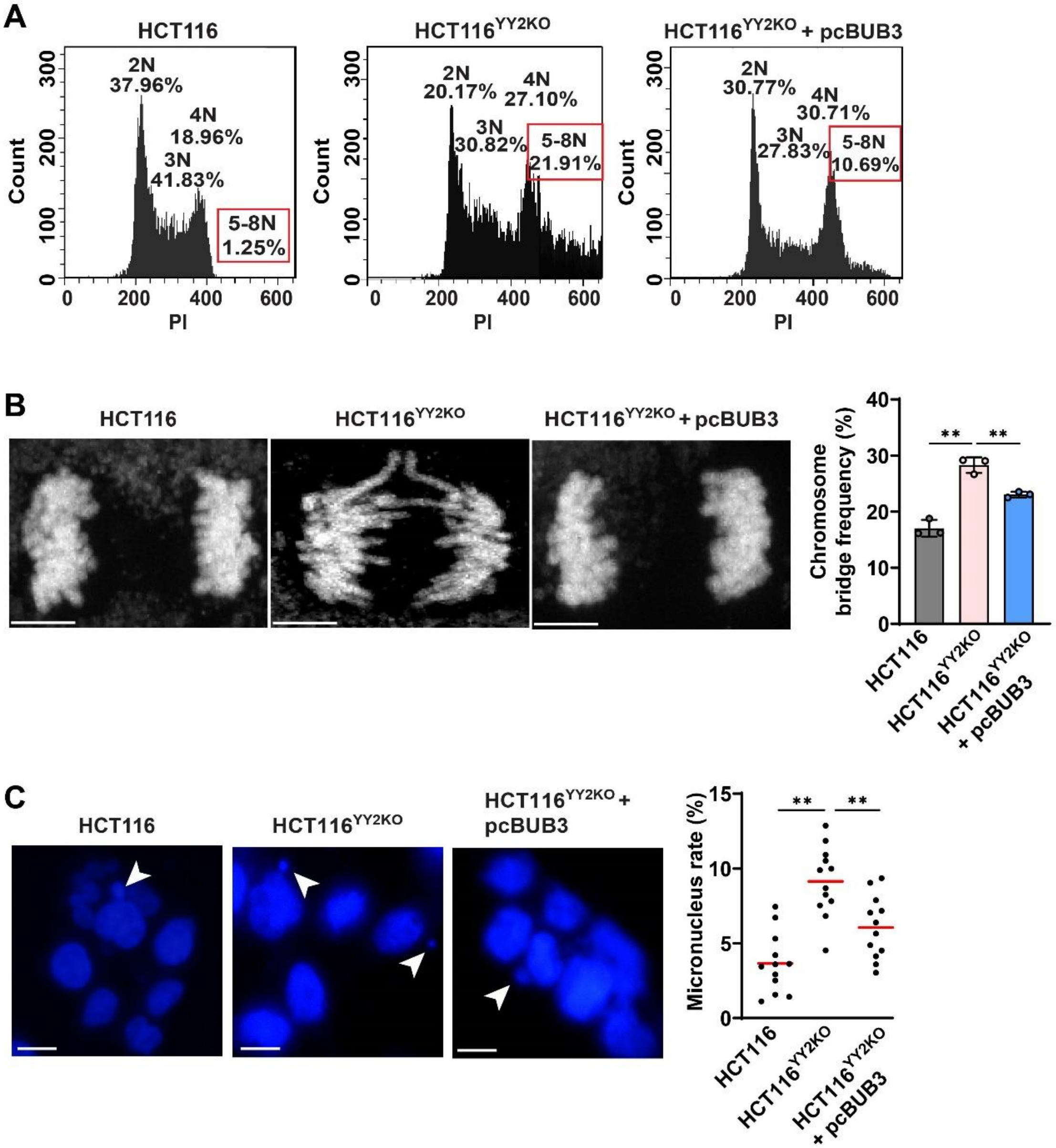


**Figure S6. Alteration in YY2 expression induces CIN.** (**A**) DNA content in *BUB3*-overexpressed HCT116$^{YY2KO}$ cells, as examined using PI staining and flow cytometry. (**B**) Chromosome bridge frequency in *BUB3*-overexpressed HCT116$^{YY2KO}$ cells. Representative images (scale bars: 5 μm) and quantification results (each dot represents chromosome bridge frequency from one independent experiment, with total 100 mitotic-cells/group) are shown. (**C**) Micronucleus rate in *BUB3*-overexpressed HCT116$^{YY2KO}$ cells. Representative images of micronuclei (indicated by arrowheads; scale bars: 20 μm) and micronucleus rate (ratio of micronuclei number to total cell number; each dot represents micronucleus rate/slide with > 100 cells/slides; four technical replicates from three independent experiments) are shown. Wild-type

HCT116 cells were used as controls. Quantification data are shown as mean ± SD of three independent experiments. $P$ values were calculated by one-way ANOVA. $^{**}P < 0.01$.

#### Supplementary Figure 7

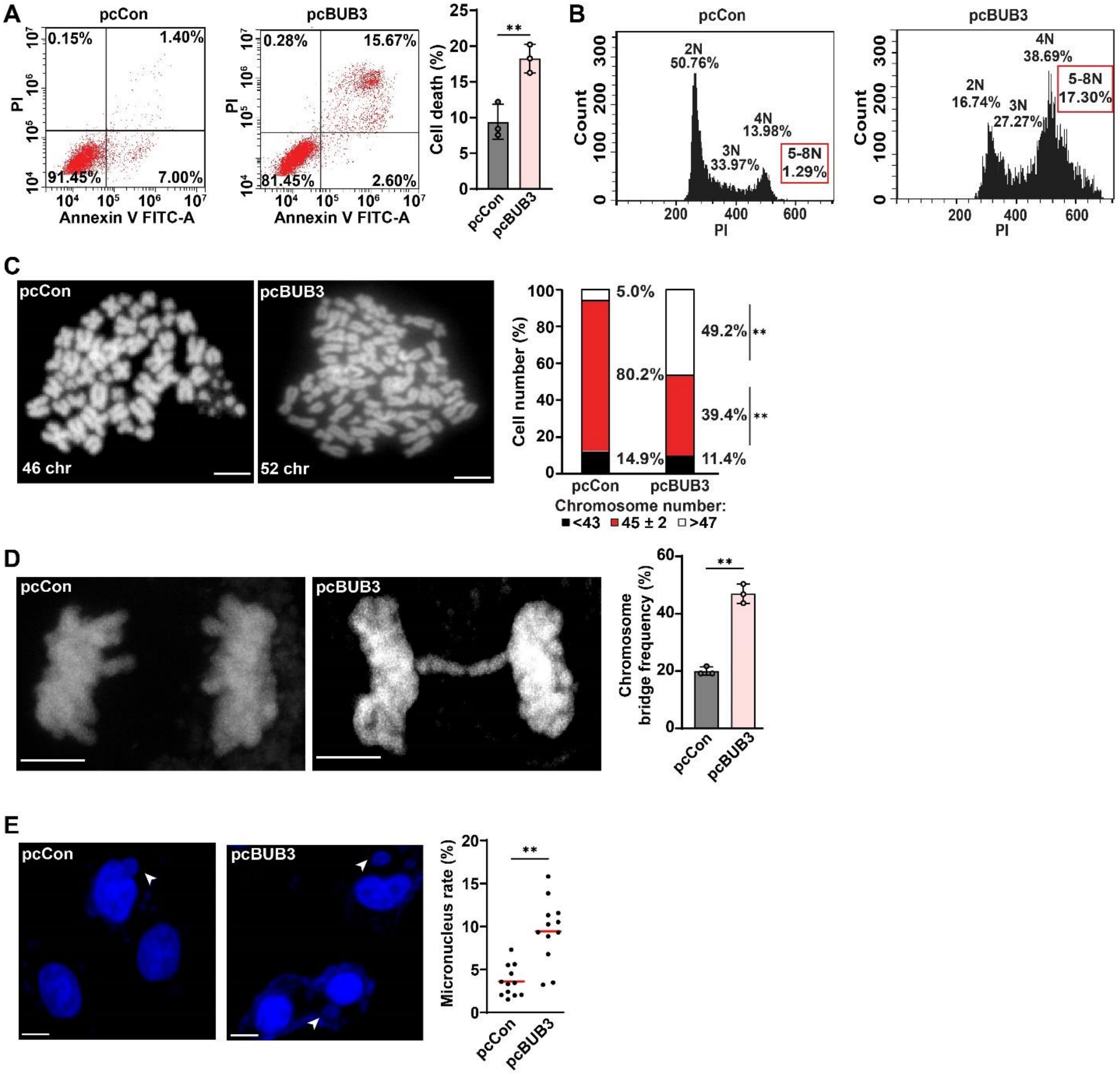


**Figure S7. *BUB3* overexpression induces CIN.** (**A**) Cell death rate of *BUB3*-overexpressed HCT116 cells, as examined using Annexin V/PI staining and flow cytometry. (**B**) DNA content in *BUB3*-overexpressed HCT116 cells, as examined using PI staining and flow cytometry. (**C**) Chromosome number per cell in *BUB3*-overexpressed HCT116 cells, as analyzed using metaphase spread. Representative images (scale bars: 10 μm) and percentage of cells with indicated chromosome number (total cells counted: 50 cells/group, pooled from three independent experiments) are shown. (**D**) Chromosome bridge frequency in *BUB3*-overexpressed HCT116 cells. Representative images (scale bars: 5 μm) and quantification results (each dot represents chromosome bridge frequency from one independent experiment, with total 100 mitotic-cells/group) are shown. (**E**) Micronucleus rate in *BUB3*-overexpressed HCT116 cells. Representative images of micronuclei (indicated by arrowheads; scale bars: 20 μm) and micronucleus rate (ratio of micronuclei number to total cell number; each dot represents micronucleus rate/slide with > 100 cells/slides; four technical replicates from three independent experiments) are shown. Cells transfected with pcCon were used as controls. Quantification data are shown as mean ± SD of three independent experiments. *P* values were calculated by one-way ANOVA. pcCon: pcEF9-Puro; $^{**}P < 0.01$.

### Supplementary Figure 8

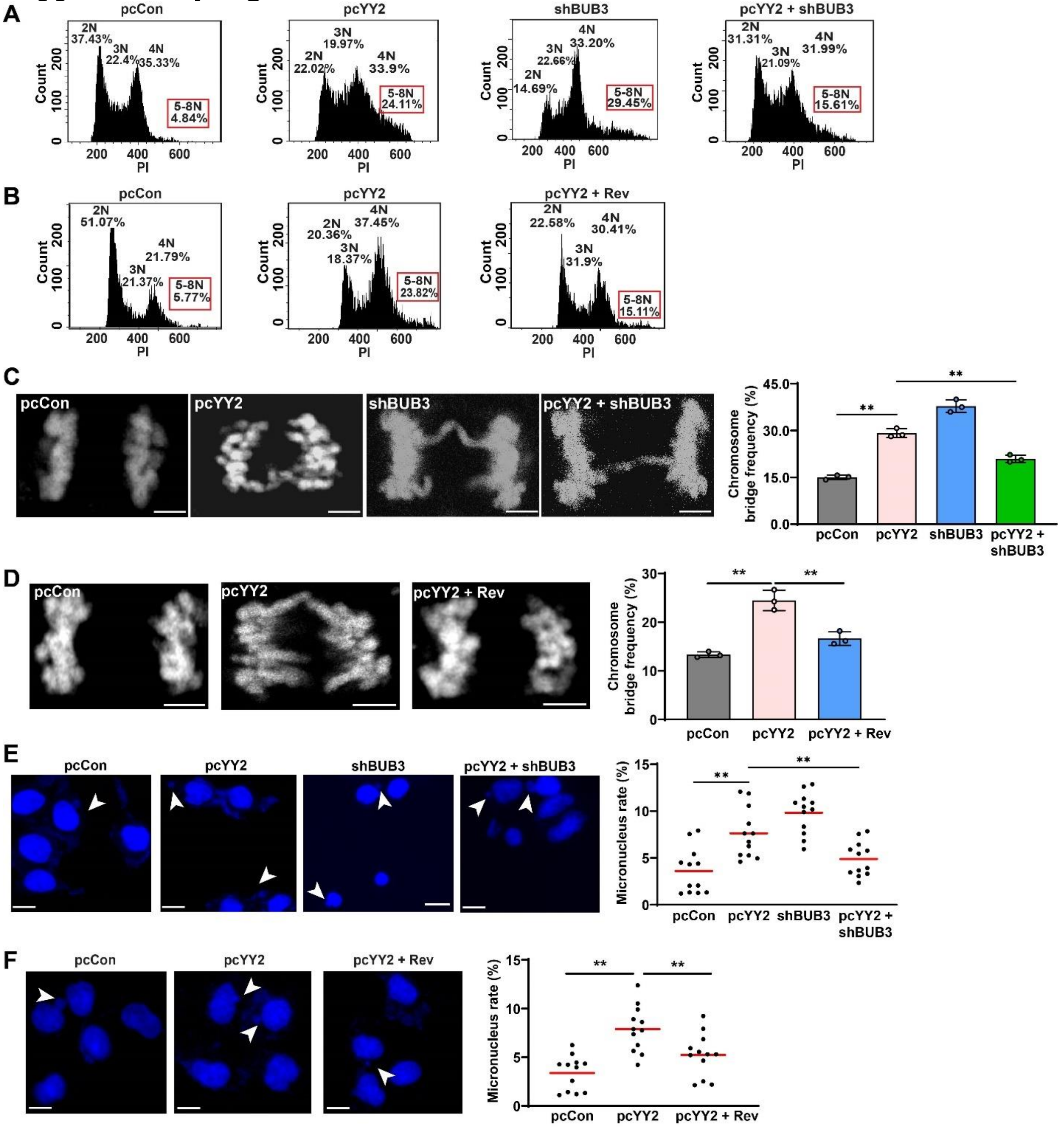


**Figure S8. *YY2* overexpression induces CIN by hyperactivating SAC.** (**A–B**) DNA content in *BUB3*-knocked down, *YY2*-overexpressed HCT116 cells (A) and *YY2*-overexpressed, reversine-treated HCT116 cells (B), as examined using PI staining and flow cytometry. (**C–D**) Chromosome bridge frequency in *BUB3*-knocked down, *YY2*-overexpressed HCT116 cells (C) and *YY2*-overexpressed, reversine-treated HCT116 cells (D), as stained using DAPI. Representative images (scale bars: 5 μm) and quantification results (each dot represents chromosome bridge frequency from one independent experiment, with total 100 mitotic-cells/group) are shown. (**E–F**) Micronucleus rate in *BUB3*-knocked down, *YY2*-overexpressed HCT116 cells (E) and *YY2*-overexpressed, reversine-treated HCT116 cells (F), as stained using DAPI. Representative images of micronuclei (indicated by arrowheads; scale bars: 20 μm) and quantification results (ratio of micronuclei number to total cell number; each dot represents micronucleus rate/slide with > 100 cells/slides; four technical replicates from three independent experiments) are shown. Cells transfected with pcCon or pcCon and shCon were used as controls. Quantification data are shown as mean ± SD. All data were obtained from three independent experiments. *P* values were calculated by one-way ANOVA**.** pcCon: pcEF9-Puro. Rev: reversine (final concentration: 0.2 μM). **$P < 0.01$.

#### Supplementary Figure 9

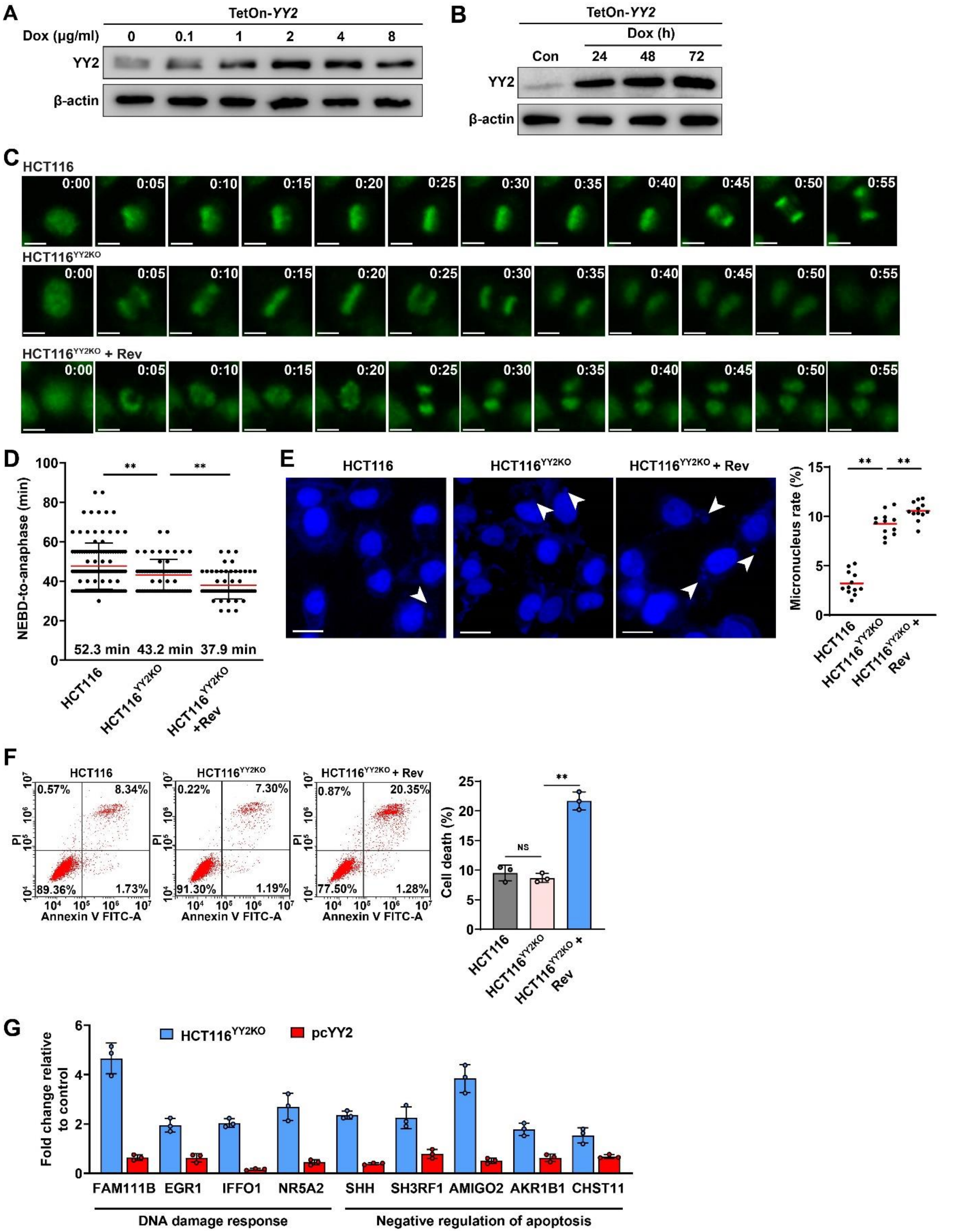


**Figure S9. SAC inhibition induces excessive CIN in *YY2*-knockout cells.** (**A–B**) YY2 protein expression level in HCT116 cells infected with TetOn-*YY2* lentivirus after treatment with indicated dose of doxycycline for 48 h (A) or with 2 μg/ml doxycycline for indicated time (B), as determined using western blotting. (**C**–**D**) Mitotic time of reversine-treated HCT116$^{YY2KO}$ cells, as determined using time-lapse microscopy. Representative images (C; scale bars: 20 μm) and scatter plot showing the time-length from NEBD to anaphase (D; n = 60, pooled from three independent experiments) are shown. (**E**) Micronucleus rate in reversine-treated HCT116$^{YY2KO}$ cells, as stained using DAPI. Representative images of micronuclei (indicated by arrowheads; scale bars: 20 μm) and quantification results (ratio of micronuclei number to total cell number; each dot represents micronucleus rate/slide with >100 cells/slides; four technical replicates from three independent experiments) are shown. (**F**) Cell death rate of reversine-treated HCT116$^{YY2KO}$ cells, as examined using Annexin V/PI staining. (**G**) Fold change of mRNA expression levels of genes related to DNA damage response and negative regulation of apoptosis in HCT116$^{YY2KO}$ cells and *YY2*-overexpressed HCT116 cells compared to wild-type HCT116 or HCT116 cells transfected with pcCon, as analyzed using qRT-PCR. β-actin was used for qRT-PCR normalization and as western blotting loading control. Quantification data are shown as mean ± SD. All data were obtained from three independent experiments. *P* values were calculated by one-way ANOVA. Rev: Reversine (final concentration: 0.2 μM); $^{**}P < 0.01$; NS: not significant.

### Supplementary Figure 10

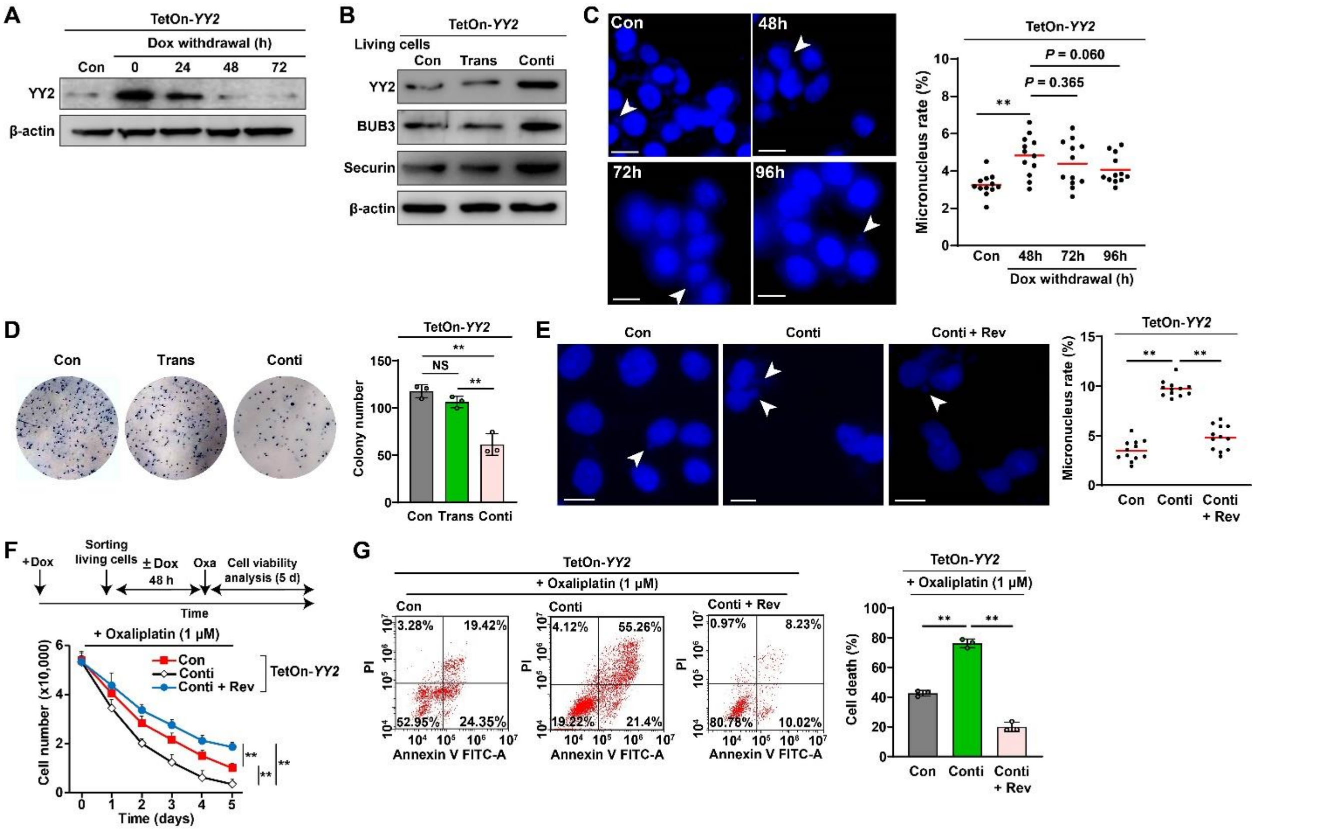


**Figure S10. YY2-transient overexpression induces heritable CIN.** (**A**) YY2 protein expression level in HCT116 cells transiently overexpressing *YY2* at indicated time-points after doxycycline-withdrawal, as determined using western blotting. (**B**) YY2, BUB3, and securin protein expression levels in HCT116 cells transiently overexpressing *YY2* (TetOn-*YY2* Trans cells), as determined using western blotting. (**C**) Micronucleus rate in HCT116 cells transiently overexpressing *YY2* at indicated time-points after doxycycline-withdrawal, as stained using DAPI. Representative images of micronuclei (indicated by arrowheads; scale bars: 20 μm) and quantification results (ratio of micronuclei number to total cell number; each dot represents micronucleus rate/slide with > 100 cells/slides; four technical replicates from three independent experiments) are shown. (**D**) Colony-formation potential of TetOn-*YY2* Trans cells at 10 days. Representative images (left) and quantification results (right; each dot represents the mean value of three technical replicates) are shown. (**E**) Micronucleus rate of reversine-treated TetOn-*YY2* Trans cells 48 h after sorting. Representative images of micronuclei (indicated by arrowheads; scale bars: 20 μm) and micronucleus rate (ratio of micronuclei number to total cell number; each dot represents micronucleus rate/slide with > 100 cells/slides; four technical replicates from three independent experiments) are shown. (**F–G**) Viability (F) and cell death rate (G) of reversine-treated TetOn-*YY2* Trans cells at indicated times and 5 days after oxaliplatin treatment, respectively. Cells infected with lentivirus generated using pTRIPZ-control (Con) were used as control. β-actin was used for western blotting loading control. Quantification data are shown as mean ± SD. All data were obtained from three independent experiments. *P* values were calculated by one-way ANOVA. TetOn-*YY2* Trans and Conti: cells infected with TetOn-*YY2* lentivirus and treated with doxycycline only before sorting or continuously, respectively; Dox: doxycycline (final concentration: 2 μg/ml); Rev: Reversine (final concentration: 0.2 μM); $^{**}P < 0.01$; NS: not significant.

#### Supplementary Figure 11

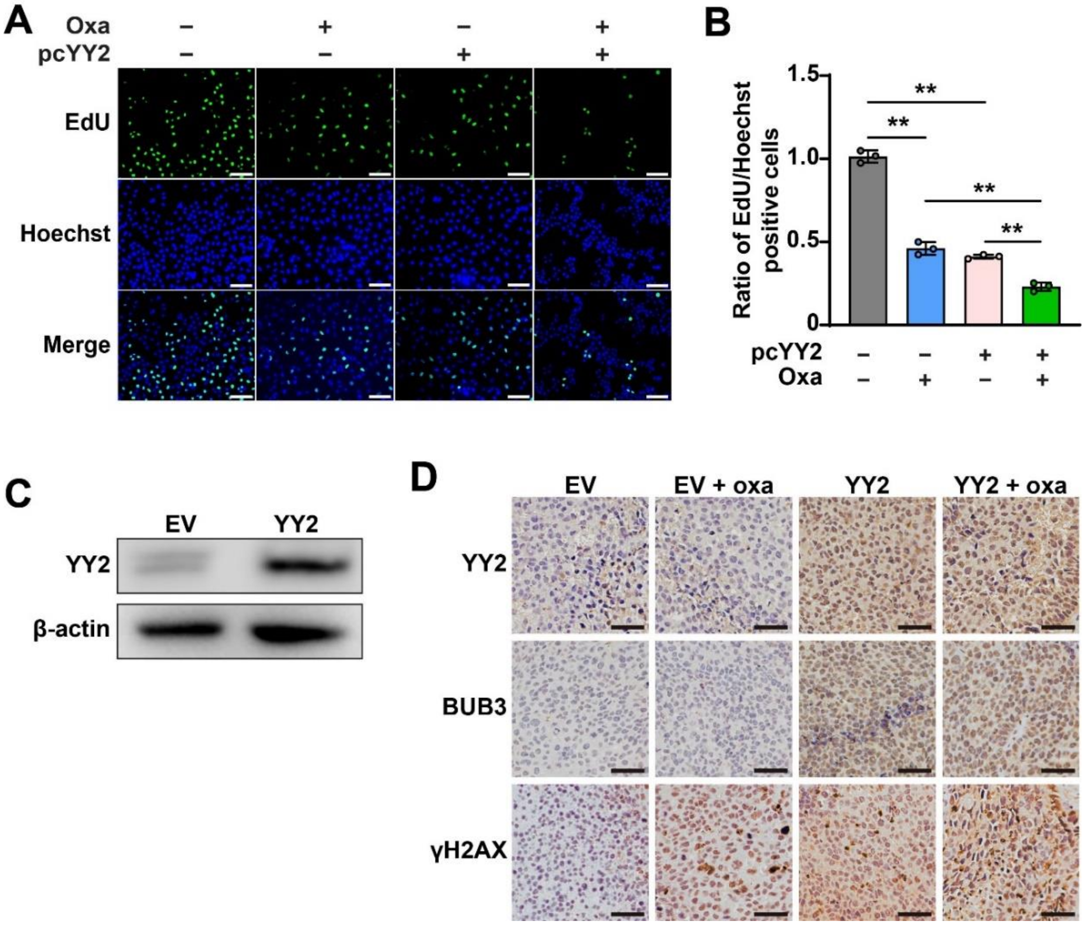


**Figure S11. YY2 enhances oxaliplatin tumor suppressive activity.** (**A–B**) Proliferation potential of oxaliplatin-treated *YY2*-overexpressed HCT116 cells as determined using EdU incorporation assay. Representative images (A, scale bars: 200 μm) and ratio of proliferative cells (B; each dot represents the mean value of three technical replicates) are shown. (**C**) YY2 protein expression level in HCT116 cells stably overexpressing *YY2* established using Lenti-YY2 virus, as determined using western blotting. (**D**) Immunohistochemistry staining images against YY2, BUB3, and γH2AX in tissue sections of xenografted tumors (scale bars: 50 μm). Cells transfected with pcCon or infected with lentivirus generated using empty lentivirus (EV) were used as control. β-actin was used as western blotting loading control. Quantification data are shown as mean ± SD. All data were obtained from three independent experiments. *P* values were calculated by one-way ANOVA**.** Oxa: Oxaliplatin; $^{**}P < 0.01$.

#### Supplementary Figure S12

**A**

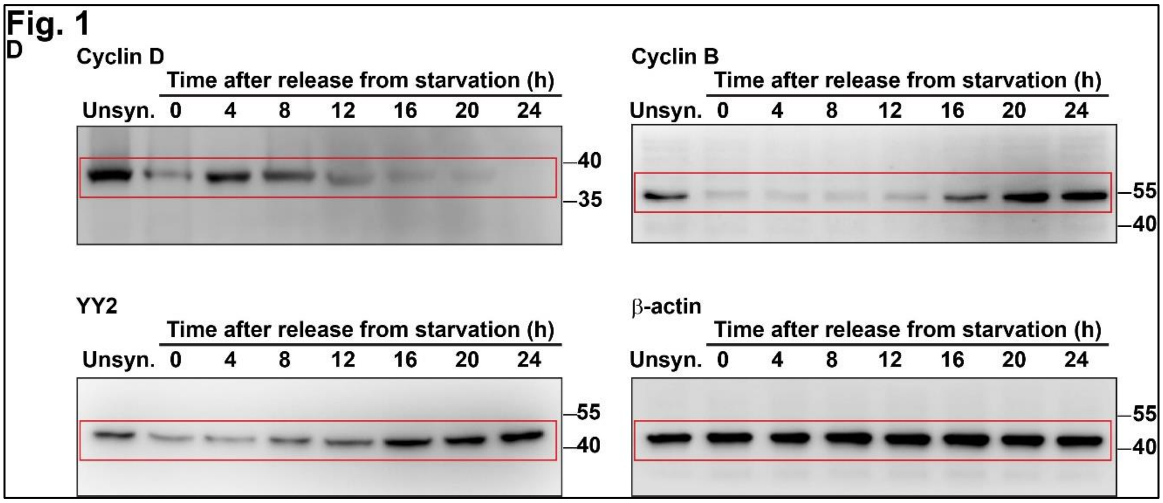


**Figure S12. Uncropped western blots with the indicated areas of selection in Figs. 1, 3, 7, and Supplementary Figs. S1, S2, S3, S4, S5, S9, S10, and S11. (continued)**

**B**

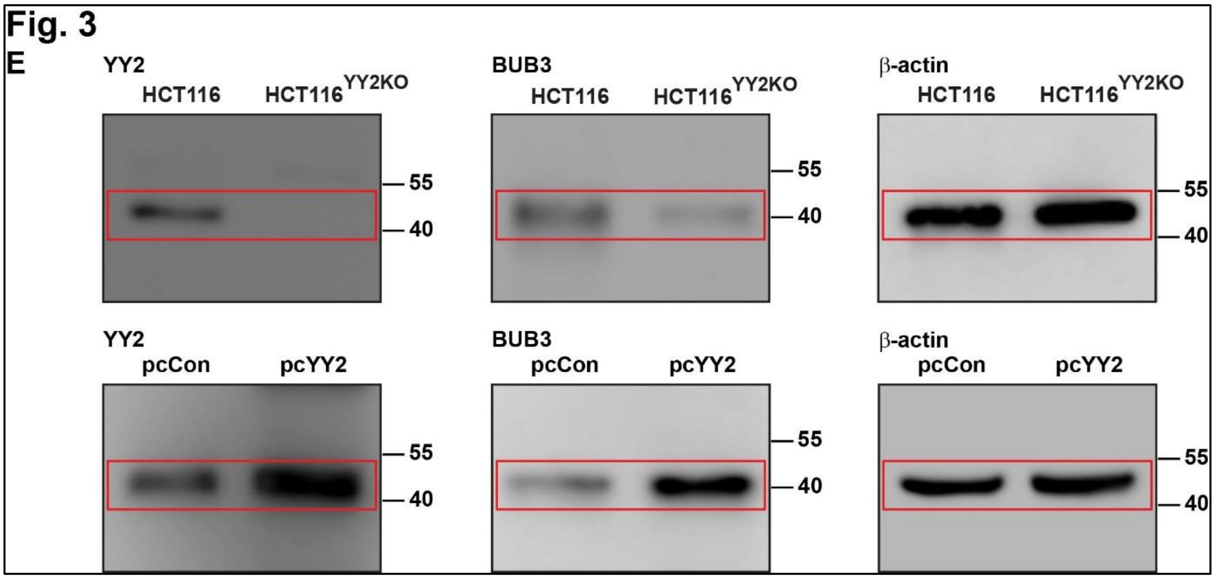


**Figure S12. Uncropped western blots with the indicated areas of selection in Figs. 1, 3, 7, and Supplementary Figs. S1, S2, S3, S4, S5, S9, S10, and S11. (continued)**

C

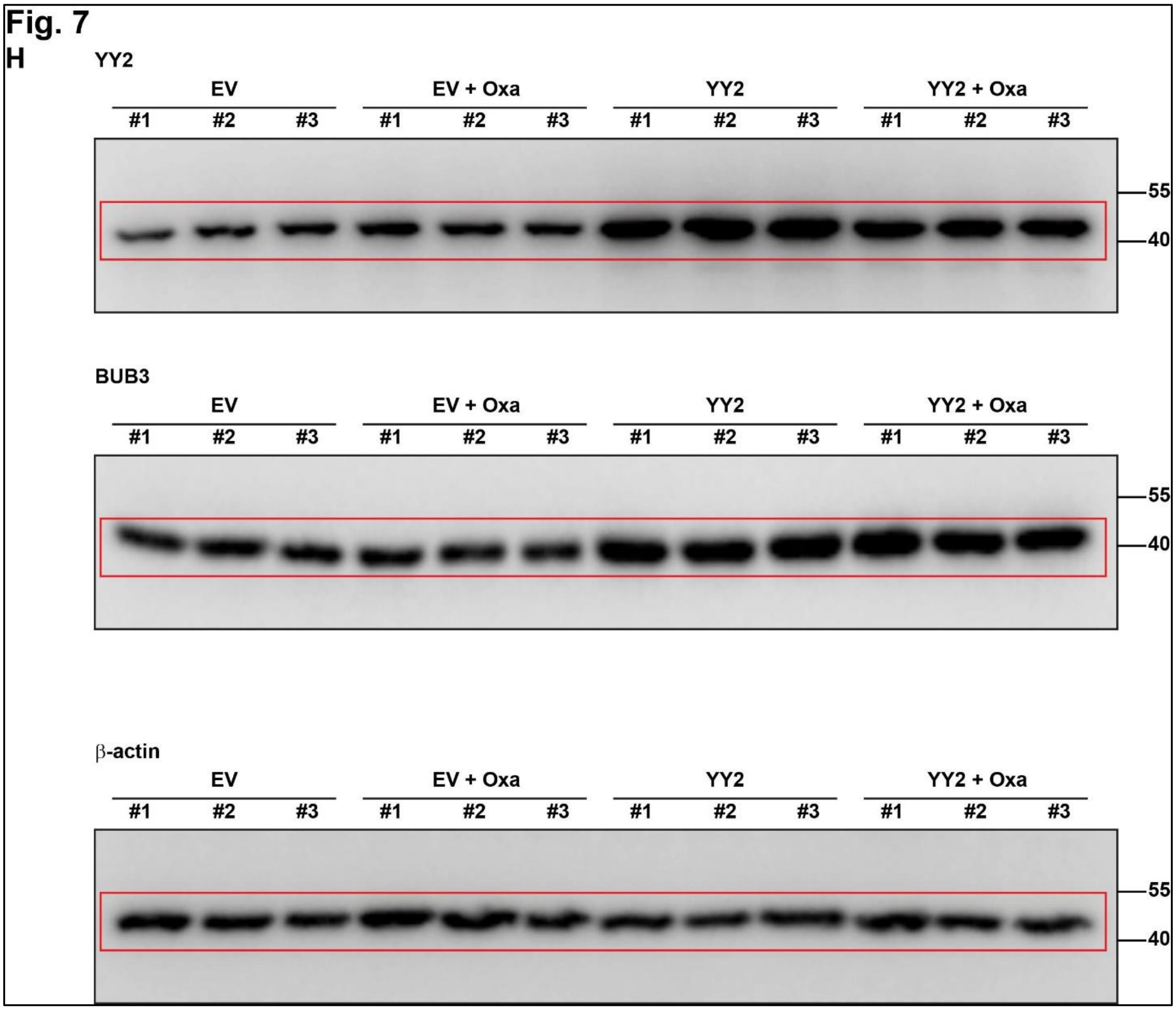


**Figure S12. Uncropped western blots with the indicated areas of selection in Figs. 1, 3, 7, and Supplementary Figs. S1, S2, S3, S4, S5, S9, S10, and S11. (continued)**

**D**

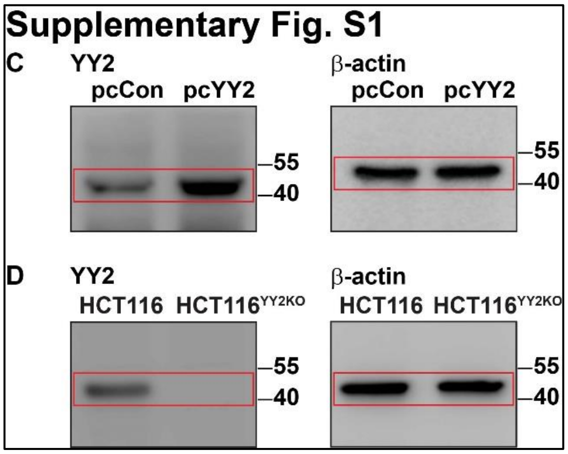


**Figure S12. Uncropped western blots with the indicated areas of selection in Figs. 1, 3, 7, and Supplementary Figs. S1, S2, S3, S4, S5, S9, S10, and S11. (continued)**

E

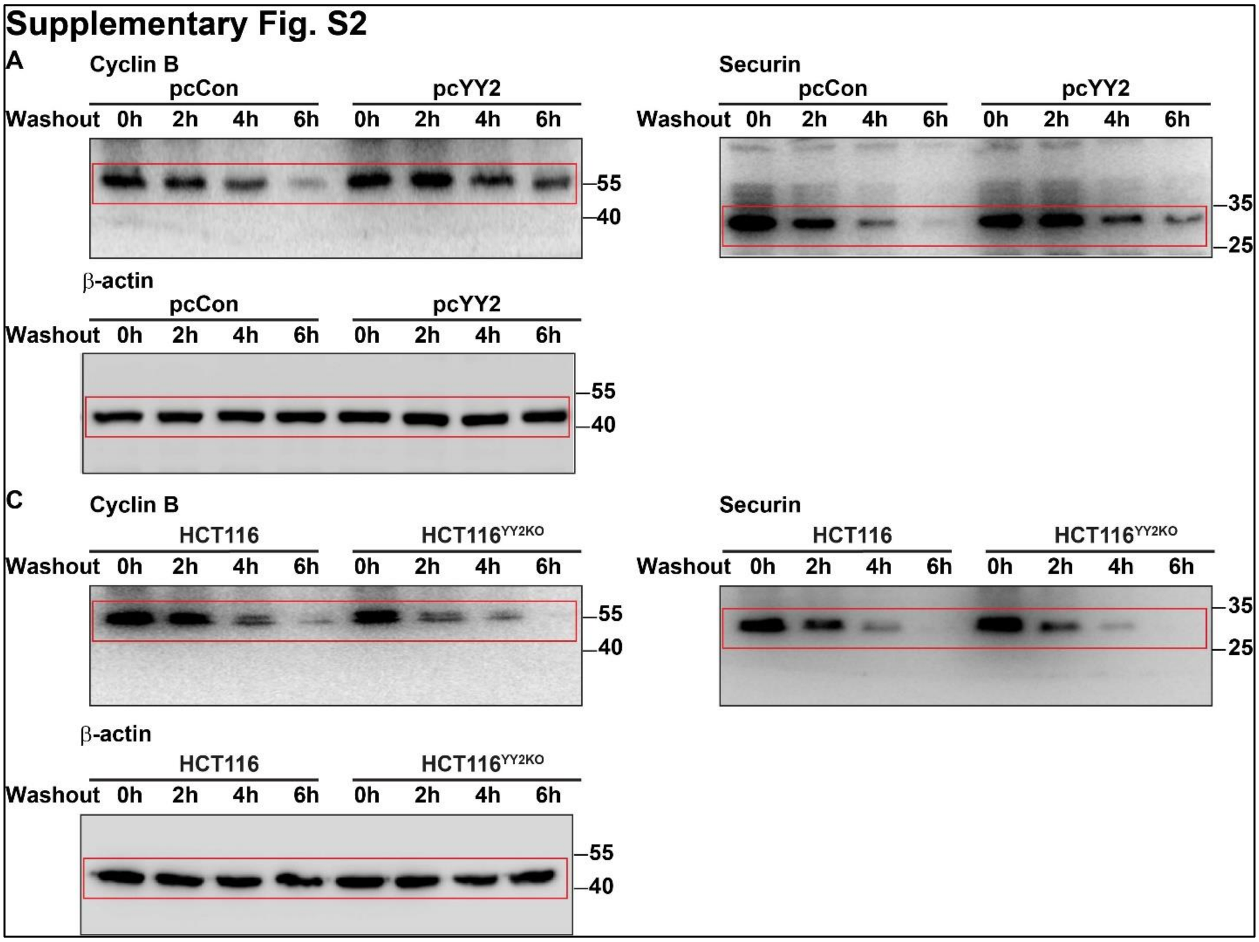


**Figure S12. Uncropped western blots with the indicated areas of selection in Figs. 1, 3, 7, and Supplementary Figs. S1, S2, S3, S4, S5, S9, S10, and S11. (continued)**

F

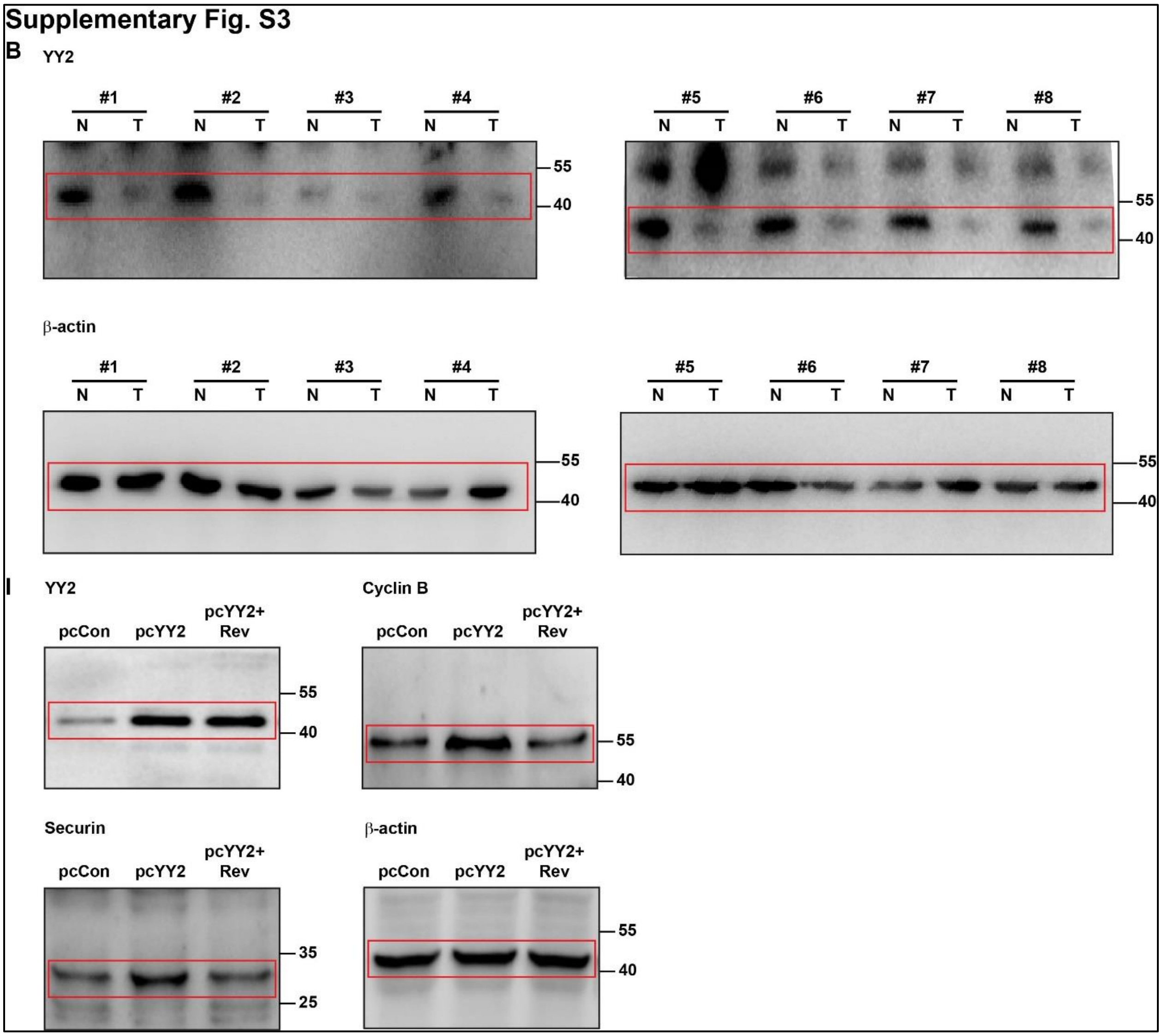


**Figure S12. Uncropped western blots with the indicated areas of selection in Figs. 1, 3, 7, and Supplementary Figs. S1, S2, S3, S4, S5, S9, S10, and S11. (continued)**

G

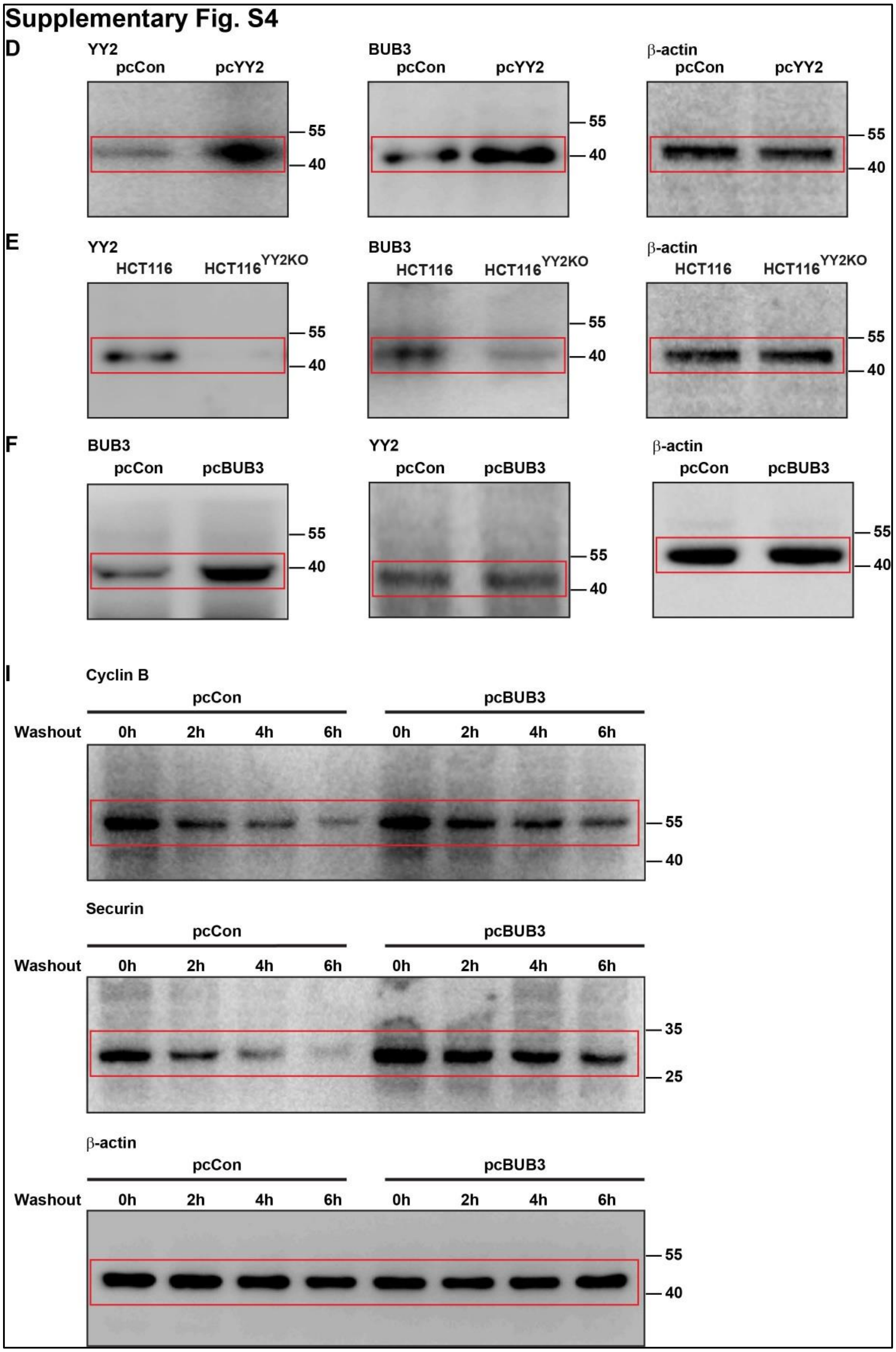


**Figure S12. Uncropped western blots with the indicated areas of selection in Figs. 1, 3, 7, and Supplementary Figs. S1, S2, S3, S4, S5, S9, S10, and S11. (continued)**

H

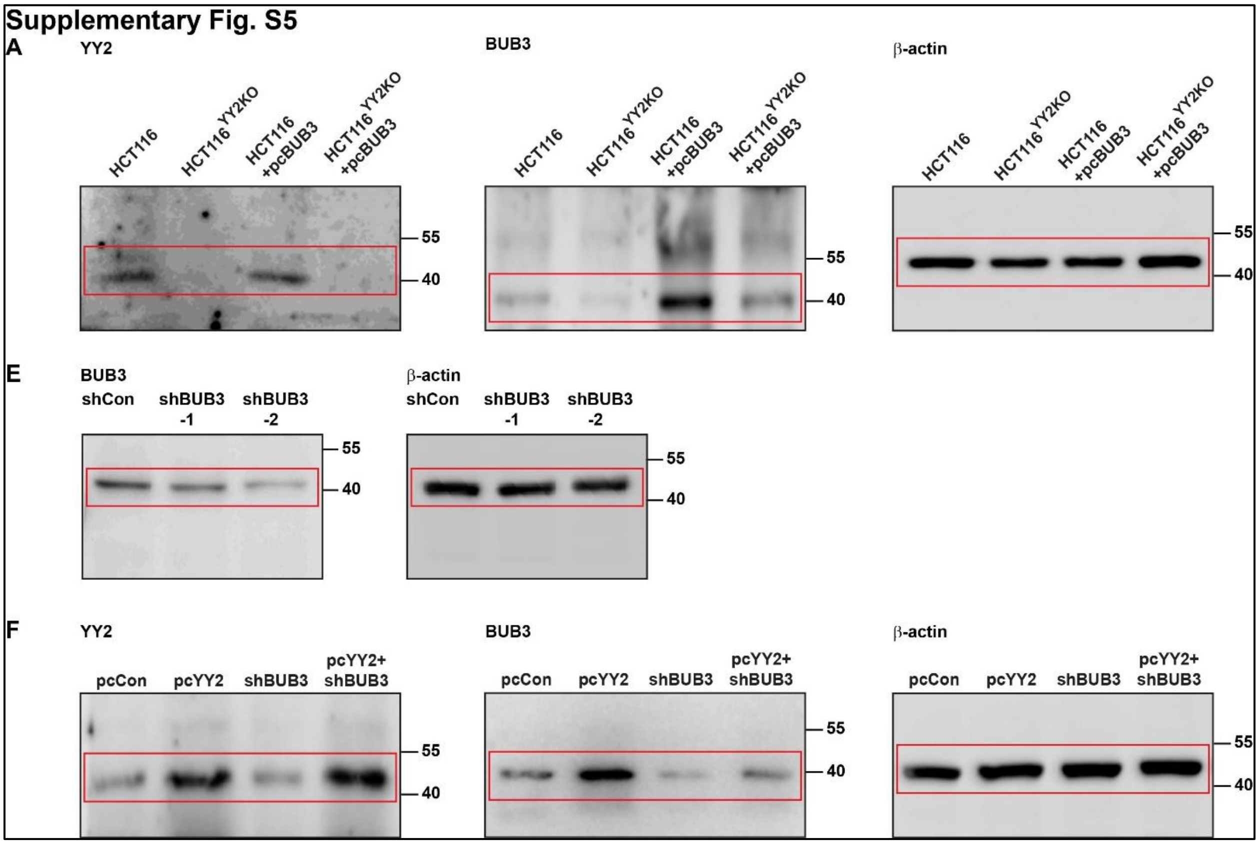


**Figure S12. Uncropped western blots with the indicated areas of selection in Figs. 1, 3, 7, and Supplementary Figs. S1, S2, S3, S4, S5, S9, S10, and S11. (continued)**

**I**

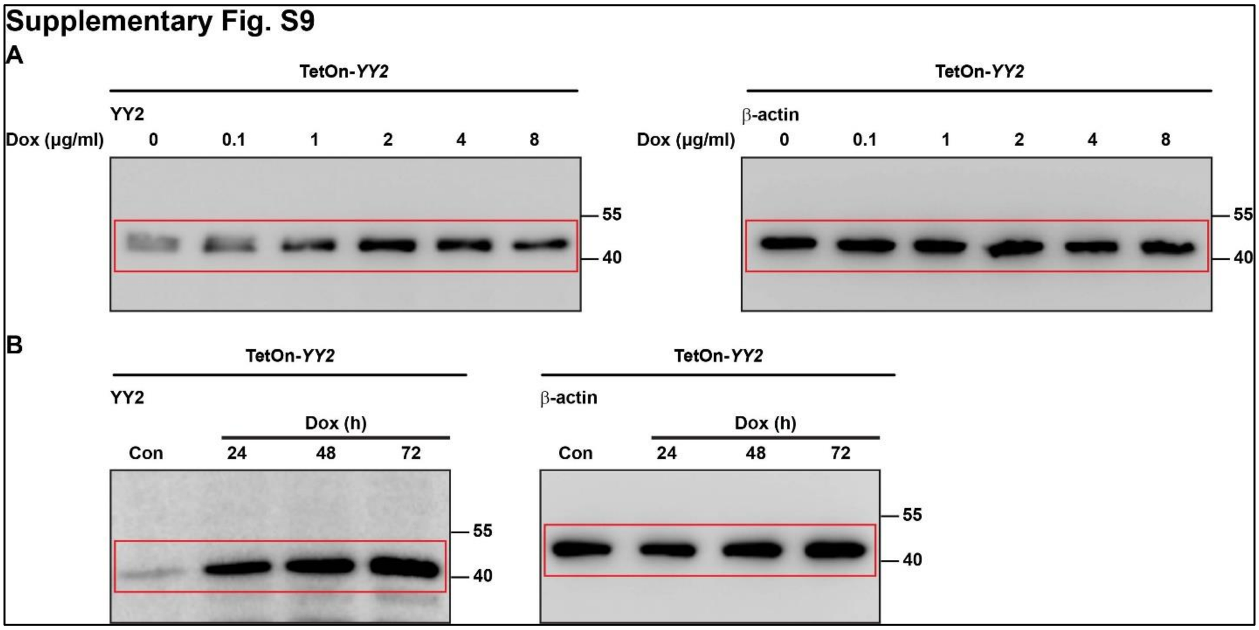


**Figure S12. Uncropped western blots with the indicated areas of selection in Figs. 1, 3, 7, and Supplementary Figs. S1, S2, S3, S4, S5, S9, S10, and S11. (continued)**

**J**

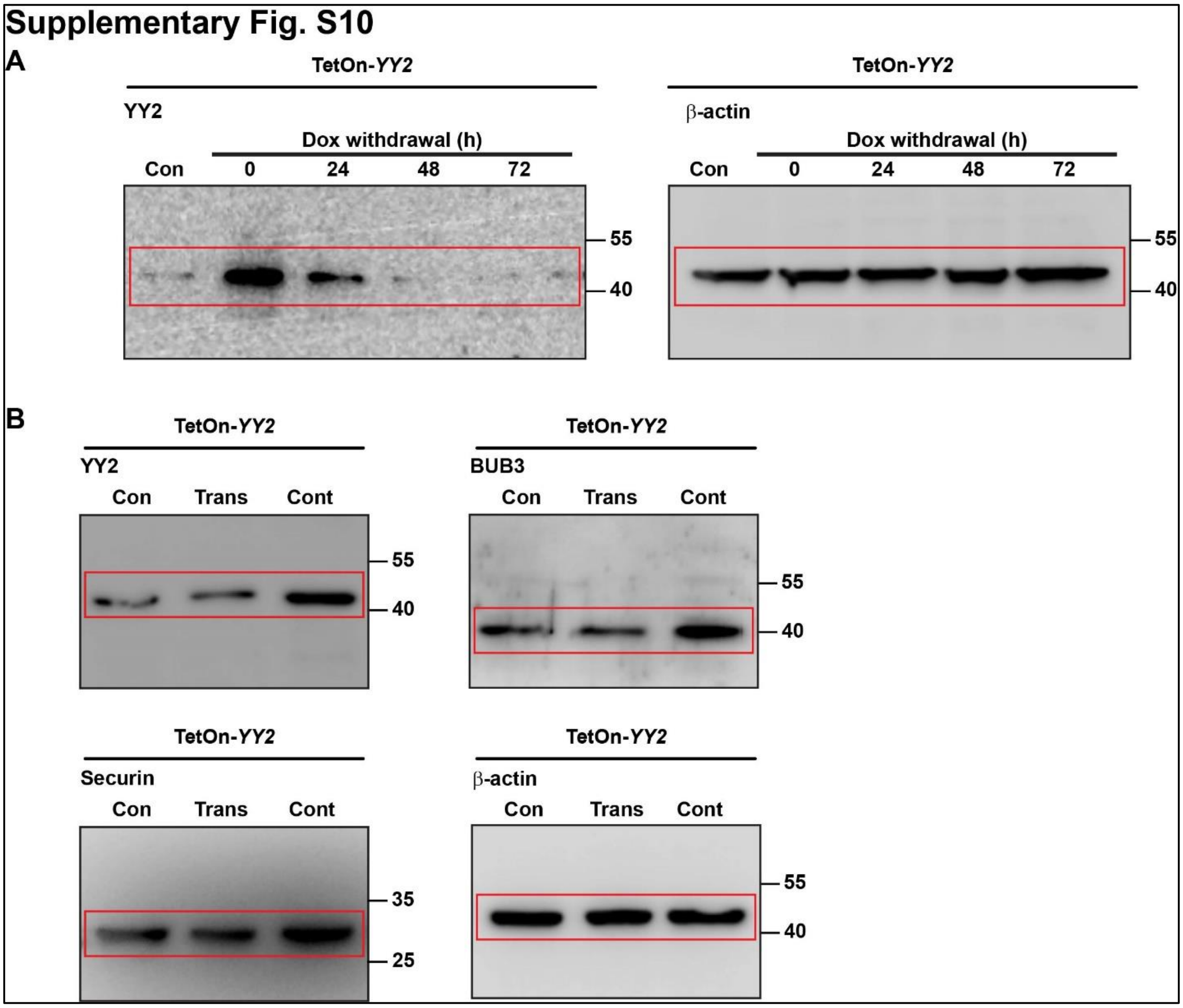


**Figure S12. Uncropped western blots with the indicated areas of selection in Figs. 1, 3, 7, and Supplementary Figs. S1, S2, S3, S4, S5, S9, S10, and S11. (continued)**

K

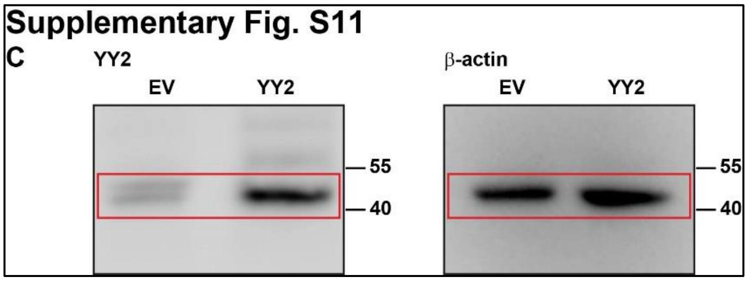


**Figure S12. Uncropped western blots with the indicated areas of selection in Figs. 1, 3, 7, and Supplementary Figs. S1, S2, S3, S4, S5, S9, S10, and S11.**

**Table S1. Primer pairs used for qRT-PCR.**

| Gene | Refseq No. | Forward primer sequence (5'-3') | Reverse primer sequence (5'-3') |
|---|---|---|---|
| YY2 | NM_206923.4 | GCAGTGGGTGAAGGCCAGGCTG | CGGTGTGGACCAGCTGGTGTCG |
| BUB3 | NM_004725.4 | AATGCTGGGACCTTCTCTCA | TCCGTAAGTCCCACACCAAC |
| TEX14 | NM_031272.5 | AGTCCCCTGTCCTGTTCAACT | GCCGCAACAAAAAGTGCTGT |
| ZFP69 | NM_001320178.2 | CAAGGCCGATGTGAAGTGGA | CTCCACGGCCTTCTTTGGAT |
| ZFP69B | NM_001369565.1 | TGACCCTTGGATCCCTGACA | TGGAAAGCTGACATCCCACT |
| ZNF93 | NM_031218.4 | ACTCTGTAACCATCCCAAACTCC | CCTCCAGGTCTCCGAAATGG |
| ZNF215 | NM_001354853.2 | CTCCTAGGGGGTTCCTTCCA | GGAGCTGGCGTTGGTAATCA |
| ZNF280C | NM_017666.5 | ATTGCCTTTAATTGGCTCTCCTC | AGCTCTTAGGTAATCTGCTCCAG |
| ZNF730 | NM_001277403.2 | CCGCCGAAGCTCCAATTTTC | GGATCTCCCAATACCCGCAG |
| ZNF878 | NM_001080404.3 | GCTCGCGCTAGTTAAGGTCT | GGCCACCGAATCCATTTCCT |
| ZNF888 | NM_001310127.2 | CTTGTAGATTCGCCCGGACC | CACATACCCAGAGTCGCCG |
| FAM111B | NM_001142703.2 | GCCCTTGAAATGCAGAATCCA | GCTGTAAACACACTACGGTCTAA |
| EGR1 | NM_001964.3 | TGATGTCCCCGCTGCAG | GTCCATGGTGGGCGAGTG |
| IFFO1 | NM_001193457.2 | CTGAACCTCCGGTTCCGCTGCTTC | GATGGGGCTGACGAAGCCGGTCTG |
| NR5A2 | NM_001276464.2 | AAGCGTTGTCCTTACTGTCG | CCCTGTCTCTCTTGTACATTGG |
| SHH | NM_000193.4 | GAAAGCAGAGAACTCGGTGG | GGAAAGTGAGGAAGTCGCTG |
| SH3RF1 | NM_020870 | GGCGCGAGAGCAAAGT | ACCTACGATCCCCAGCAAAC |
| AMIGO2 | NM_001370299.1 | GCACGAAAGGAACCATTGAT | CCTTCATGGAAACCCATTTG |
| AKR1B1 | NM_001628.4 | TTTTCCCATTGGATGAGTCGG | CCTGGAGATGGTTGAAGTTGG |
| CHST11 | NM_001173982.2 | AAACGCCAGCGGAAGAA | GGGATGGCAGAGTGAGTAGA |
| β-Actin | NM_001101.3 | CGAGCGCGGCTACAGCTT | TCCTTAATGTCACGCACGATTT |

**Table S2. Antibodies used for western blotting, immunohistochemistry, and ChIP assay.**

| Antibody | Product No. | RRID | Maker | Experiment | Dilution |
|---|---|---|---|---|---|
| anti-YY2 | sc-374455 | AB_10988247 | Santa Cruz Biotechnology | Western blotting<br>ChIP<br>IHC | 1/1,000<br>30 μg/mL cell lysate<br>1/100 |
| anti-cyclin D1 | sc-8396 | AB_627344 | Santa Cruz Biotechnology | Western blotting | 1/1,000 |
| anti-cyclin B1 | sc-245 | AB_627338 | Santa Cruz Biotechnology | Western blotting | 1/1,000 |
| anti-BUB3 | 27073-1-AP | AB_2880743 | Proteintech | Western blotting<br>IHC | 1/5,000<br>1/100 |
| anti-securin | sc-56207 | AB_785382 | Santa Cruz Biotechnology | Western blotting | 1/500 |
| anti-γH2AX | GB111841 | | Servicebio | IHC | 1/100 |
| anti-β-actin | 66009-1-Ig | AB_2687938 | Proteintech | Western blotting | 1/50,000 |
| Goat Anti-Rabbit IgG | ZB2301 | AB_2747412 | ZSGB-BIO | Western blotting | 1/10,000 |
| Goat Anti-Mouse IgG | ZB2305 | AB_2747415 | ZSGB-BIO | Western blotting | 1/10,000 |

**Video Legends**

**Video S1.** Time-lapse video of control HCT116 cells related to Fig. 1. The display rate is one frame every 200 millisecond. Still images of this video are shown in Fig. 1E.

**Video S2.** Time-lapse video of *YY2*-overexpressed HCT116 cells related to Fig. 1. The display rate is one frame every 200 millisecond. Still images of this video are shown in Fig. 1E.

**Video S3.** Time-lapse video of wild-type HCT116 cells related to Fig. 1. The display rate is one frame every 200 millisecond. Still images of this video are shown in Fig. 1F.

**Video S4.** Time-lapse video of HCT116$^{YY2KO}$ cells related to Fig. 1. The display rate is one frame every 200 millisecond. Still images of this video are shown in Fig. 1F.

**Video S5.** Time-lapse video of *YY2*-overexpressed, reversine-treated HCT116 cells related to Fig. 2. The display rate is one frame every 200 millisecond. Still images of this video are shown in Fig. 2C.

**Video S6.** Time-lapse video of *BUB3*-overexpressed HCT116 cells related to Supplementary Fig. S4. The display rate is one frame every 200 millisecond. Still images of this video are shown in Supplementary Fig. S4I.

**Video S7.** Time-lapse video of *BUB3*-overexpressing HCT116$^{YY2KO}$ cells related to Supplementary Fig. S5. The display rate is one frame every 200 millisecond. Still images of this video are shown in Supplementary Fig. S5B.

**Video S8.** Time-lapse video of *BUB3* knocked-down HCT116 cells related to Supplementary Fig. S5. The display rate is one frame every 200 millisecond. Still images of this video are shown in Supplementary Fig. S5G.

**Video S9.** Time-lapse video of *BUB3* knocked-down, *YY2*-overexpressed HCT116 cells related to Supplementary Fig. S5. The display rate is one frame every 200 millisecond. Still images of this video are shown in Supplementary Fig. S5G.

**Video S10.** Time-lapse video of reversine-treated HCT116$^{YY2KO}$ cells related to Supplementary Fig. S9. The display rate is one frame every 200 millisecond. Still images of this video are shown in Supplementary Fig. S9C.
